# Tryptophan Restores Recovery Competence Following Starvation in the Absence of Lysosomal Amino Acid Sensing

**DOI:** 10.64898/2026.10.27.762351

**Authors:** John T. Murphy, Ryan Chang, Ziang Bi, Siana Kapoor, Haoying Zhang, David Seamons, Agastya Diwan, Micklaus Garcia, Moydul Islam, Kenji Rowel Q. Lim, Daniel L. Schneider, Brian M. Egan, David Rawnsley, Xiucui Ma, Kerry Kornfeld, Abhinav Diwan

**Author notes:** **Corresponding Author:** Abhinav Diwan, M.D.; Professor of Medicine; or Kerry Kornfeld, M.D. Ph.D.; Professor of Developmental Biology., Washington University School of Medicine, Division of Cardiology, 660 S. Euclid, CSRB 822 NTA, St. Louis, MO 63110.

## Abstract

How organisms recover from starvation to nutrient-replete physiology remains poorly understood. In *C. elegans*, we previously observed that transcription factor *hlh-30* or a combination of glucose and linoleic acid in its absence are essential for survival with refeeding following starvation, i.e. recovery competence. A forward genetic screen to evaluate mechanisms identified *depd-5*, which encodes a GATOR1 subunit, loss of which is expected to impair amino acid sensing with unrestrained *raga-1* and mTOR activation. Surprisingly, glucose refeeding reduced recovery competence in *depd-5(lf*, loss-of-function) animals, which was rescued by amino acids, specifically tryptophan. Increasing tryptophan levels in *depd-5(lf)* worms through supplementation, reduced tryptophan catabolism with *tdo-2* ablation and prolonged starvation restored recovery competence. *raga-1(gf*, gain-of-function) recapitulated whereas *raga-1(lf)* rescued the impaired recovery competence in starved *depd-5(lf)* worms. An analogous *DEPDC5* mutation in mammalian cells impaired starvation-induced mTOR inactivation. We conclude that tryptophan counters unrestrained mTOR activity to restore recovery competence following starvation.

## Introduction

The ability to successfully transition from starvation to recovery upon the reintroduction of nutrients is essential for resumption of growth, development and function. Observations from clinical and biological contexts challenge the assumption that restoring the food supply is sufficient to correct the deficits and lead to a successful transition. Indeed, refeeding after prolonged fasting has adverse clinical consequences, which can be modified with appropriately tailored clinical protocols.^1^ The use of graded or progressive reintroduction of nutrients increases the likelihood of a successful transition though this treatment is based on empirical observation rather than an understanding of the underlying physiology.^2^ Similarly, restoring blood flow is the gold standard treatment for myocardial infarction and stroke resulting from occluded vasculature; but this often results in reperfusion injury indicating that simple restoration of blood supply does not guarantee recovery.^3^ Understanding the mechanisms that govern the transition from starvation to recovery may offer therapeutic opportunities to improve outcomes in these conditions.

Despite significant advances in our understanding of starvation and fasting physiology,^4,5^ successful recovery after refeeding is assumed to occur via reversal of this physiology, where recovery is predicated upon nutrient restoration. Observations of long-lived *C. elegans* mutants suggests that starvation is an actively regulated rather than a passive state.^6,7^ Signaling pathways that reduce the rate of nutrient depletion and coordinate adaptive responses increase the likelihood of survival during nutrient limitation.^8^ Consistent with this view, our studies have shown that HLH-30, the worm ortholog for MiT/TFE family of master transcriptional regulators of autophagy and lysosome biogenesis, is essential to maintain survival during starvation.^9^ Importantly, not all systems are shut down, the organism actively preserves the capacity for rapid reactivation upon nutrient restoration. For example, RNA Polymerase II is positioned on promoters of growth-associated genes allowing transcription to resume rapidly when nutrients become available.^10^ Extensive regulation of the starvation state suggests that recovery is also orchestrated through actively regulated processes during both starvation and following restoration of nutrient supply. Our previous work established this as ‘recovery competence’, a genetically regulated physiological process distinct from starvation survival.^9^ HLH-30 was not only essential in promoting survival during starvation, but was also required for recovery under conditions where standard nutrient sources were restored. Glucose and linoleic acid bypassed the requirement for HLH-30, demonstrating that specific nutrient-responsive pathways can restore recovery competence independently of canonical HLH-30 function.^9^

To identify the molecular mechanism responsible, we performed an unbiased forward genetic screen to uncover enhancers of glucose and linoleic acid-mediated restoration of recovery competence in starved *hlh-30* null animals. This screen identified *depd-5*, encoding a component of the conserved GATOR1 complex that was proposed to be essential for ‘sensing’ amino acid availability.^11^ In this paradigm, amino acid starvation triggers GATOR1’s GAP (GTPase activating protein sense) activity to inactivate RAGA and inhibit mTOR activity.^11^ Characterization of *depd-5* mutant worms revealed that nutrient restoration alone is insufficient to ensure successful recovery with provision of glucose and other energy sources. Surprisingly, recovery required amino acids, specifically tryptophan in the setting of *depd-5* loss, indicating that tryptophan availability is sensed through yet undiscovered pathways when canonical lysosomal amino acid signaling is impaired and restores recovery competence when an energy source is available.

## Results

### A Genetic Screen for Mutations that Regulate Recovery Competence Following Starvation in *C. elegans* Identified *depd-5*

*hlh-30(lf)* mutants display a defect in starvation recovery compared to wild type (WT) animals,^9^ indicating that *hlh-30* is necessary for starvation recovery, a capacity we define as recovery competence. We discovered that *hlh-30(lf)* mutants display improved starvation recovery when they are exposed to CeMM (*C. elegans* maintenance medium^12^) prior to culturing on live bacteria; the key ingredients in CeMM that promote improved starvation recovery are glucose and linoleic acid, which act in a time-dependent manner.^9^ To discover new genes required for recovery competence, we used ethyl methanesulfonate (EMS) to mutagenize *hlh-30(tm1978)* mutants and performed an F3 clonal forward genetic screen (**Figure 1A**). To assess recovery competence, we starved worms for 33 hours, refed with glucose and linoleic acid in CeMM minimal media **(Table S1)** for 15 hours, transferred to agar dishes layered with *E. coli* OP50, and assessed survival after 48 hours (**Figure 1B**).

**Figure 1.**
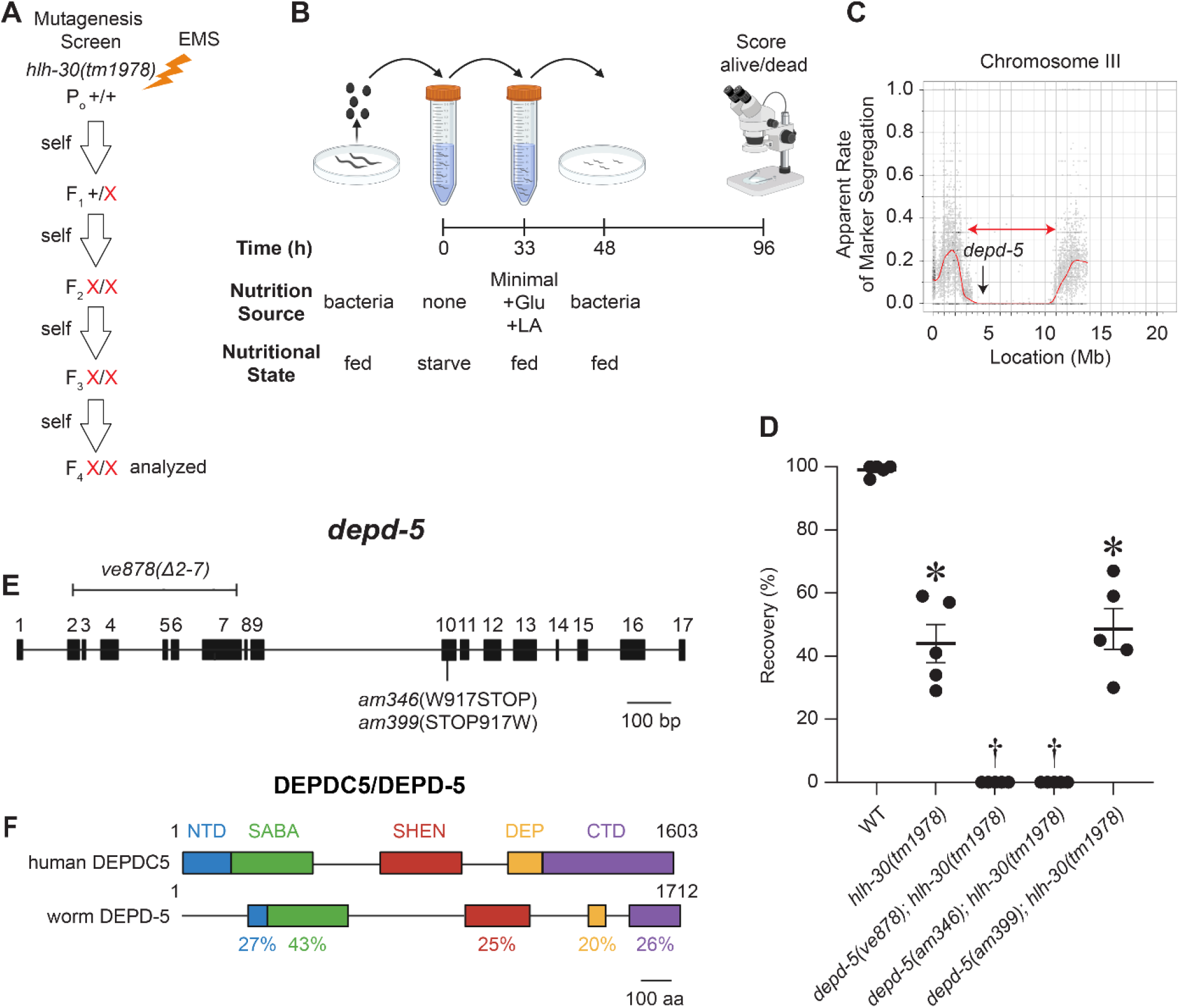
A forward genetic screen leads to the discovery of *depd-5*, a regulator of recovery competence following starvation. **(A)** Schematic depicting the EMS mutagenesis screen. The P_0_ generation of *hlh-30(tm1978)* mutants was treated with the mutagen ethyl methanesulfonate (EMS). Wild-type alleles are shown as +, while new mutant alleles are shown as ‘X’. The F_4_ generation was screened for animals that failed to recover following the workflow described in B. **(B)** Diagram of the experimental workflow for the assessment of starvation recovery (%). Gravid adults are fed bacteria, treated with alkaline hypochlorite, and their progeny hatch in a nutrient-free M9 salt solution (time 0 – 33 hours). Larvae are first refed by incubating in the chemically defined liquid media, *C. elegans* Maintenance Media (CeMM) (time 33-48 hours). Larvae are then cultured on dishes seeded with *E. coli* OP50 (time 48-96 hours). Recovery (%) is defined as the number of animals alive at 96 hours divided by the number of animals alive at 48 hours. **(C)** Mapping plot depicting Hawaiian SNPs generated from pooled sequencing of recombinant DNA. The horizontal axis shows the physical map of chromosome III (megabases), and the vertical axis shows the rate of Hawaiian SNP marker segregation. The red horizontal arrow indicates the mapping region with a very low frequency of Hawaiian SNP markers; the black vertical arrow marks the approximate position of a candidate nonsense mutation in *depd-5*. **(D)** The horizontal axis shows the genotypes, and the vertical axis shows percent recovery as defined in B. Bars and whiskers represent mean ± SEM (five independent experiments). The *ve878; tm1978* and *am346; tm1978* mutants displayed 0% recovery and therefore had no variance. These groups were not included in the statistical analysis. Statistical significance among the remaining groups was determined by ordinary one-way ANOVA followed by Tukey’s multiple-comparisons test. \**P* < 0.05 versus wild type. †0% recovery in all five independent experiments. **(E)** The gene structure of *depd-5* depicts exons as boxes and introns as connecting lines. The *ve878* deletion is shown above the gene structure. The *am346* nonsense mutation (W917STOP) and the *am399* reversion mutation (STOP917W) are shown below. Scale bar = 100 base pairs. **(F)** Comparison of amino acid domains between human DEPDC5 and *C. elegans* DEPD-5 proteins. Conserved domains (NTD, SABA, SHEN, DEP and CTD) are depicted as boxes connected by lines. Percent amino acid identify between corresponding domains was determined by pairwise sequence alignment using EMBL-EBI EMBOSS Needle and is shown below the *C. elegans* DEPD-5 protein. Numbers indicate the first and last amino acid of each protein. Scale bar = 100 amino acids.

We isolated the *am346* mutant strain based on reduced recovery competence following starvation (**Figure 1D**). To identify the molecular lesion that causes the *am346* phenotype, we used a positional cloning strategy based on single nucleotide polymorphism (SNP) mapping and whole-genome sequencing.^13^ The results showed that the *am346* mutation maps to a region of chromosome III (**Figure 1C, S1**). We identified candidate EMS mutations in this region that are predicted to cause missense and nonsense mutations. Two genes contained nonsense mutations: *C09E7.10* and *T08A11.1* **(Table S2, S3**). To ascertain the causative gene, we reverted the W917STOP mutation in *T08A11.1* using CRISPR targeting, generating the *am399* allele (**Figure 1D, E**). We starved the control strain, *am346; hlh-30(tm1978)* and *am399; hlh-30(tm1978)* mutants for 33 hours at the L1 larval stage and exposed these animals to CeMM minimal media supplemented with glucose and linoleic acid for 15 hours, recapitulating the conditions of the screen. The mutant and control strains displayed no lethality immediately following treatment with glucose and linoleic acid in CeMM minimal (**Figure S2**). When these animals were transferred to NGM agar with *E. coli* and scored for survival after 48 hours, the *am346; hlh-30(tm1978)* demonstrated a complete lack of recovery, indicating impaired recovery competence as compared with wild type (WT, **Figure 1D**). Importantly, the reverted *am399; hlh-30(tm1978)* strain displayed restored recovery competence to levels observed in the parental *hlh-30(tm1978)* strain, suggesting the nonsense (W917STOP) mutation in *T08A11.1* is causative (**Figure 1D**). Moreover, we obtained the independently derived *T08A11.1* mutation *ve878* (**Figure 1E)** and found that it also reduced starvation recovery in the *hlh-30(tm1978)* background (**Figure 1D**). These data establish *am346* as a loss-of-function allele of *T08A11.1*. We also employed CRISPR gene editing in the *am346* strain to revert the R94STOP mutation in *C09E7.10* (**Table S3**) to the wild-type sequence; rescue was not observed (data not shown), indicating this mutation does not cause the phenotype.

We subjected the predicted *T08A11.1* protein to Basic Local Alignment Search Tool (BLAST) analysis against the human proteome and identified homology with human DEP domain containing 5, DEPDC5^14^ (**Figure 1F, S3**). In mammals, DEPDC5 functions with NPRL2 and NPRL3 in the trimeric GAP Activity Towards Rags 1 (GATOR1) complex, a conserved negative regulator of mTORC1 signaling.^11^ Loss-of-function mutations in human DEPDC5 are associated with focal epilepsy.^15,16^ We therefore named the *C. elegans T08A11.1* gene *depd-5*, and designated *am346* as an allele of *depd-5*. These findings identified *depd-5* as a previously unrecognized gene that promotes recovery competence after starvation.

### Recovery After Starvation in *depd-5* Mutant Worms is Impaired by Glucose and Improved by Tryptophan

*depd-5(am346); hlh-30(tm1978)* mutants failed to recover from starvation on CeMM minimal medium containing glucose and linoleic acid (**Figure 1D**). To evaluate the phenotype of *depd-5(am346)* in an otherwise wild-type background (with intact *hlh-30*), we used a genetic cross to separate *am346* from the *hlh-30(tm1978)* mutation. In subsequent experiments, *depd-5(am346)* worms were starved for 24 hours and exposed to glucose and linoleic acid in CeMM minimal medium for 24 hours (**Figure S4A**). These animals did not display lethality at this stage (**Figure S4B**). They were then cultured on *E. coli* OP50 agar dishes for 48 hours and assessed for survival. Glucose in combination with linoleic acid significantly reduced recovery in *depd-5*(*am346*) mutants compared to wild type (**Figure 2A**). Because glucose supports recovery after starvation in both WT and *hlh-30(tm1978)* mutants,^9^ its detrimental effect was unexpected. Therefore, we tested the effect of other nutrient classes on starvation recovery. Administration of nucleic acids did not enhance recovery from starvation (**Figure S4C**). By contrast, administration of all amino acids restored recovery competence in *depd-5(am346)* mutants (**Figure 2A**). This was also unexpected as DEPDC5, the mammalian ortholog of DEPD-5, is a component of the GATOR1 complex which has been ascribed a key role in amino acid sensing, and mammalian cells deficient in DEPDC5 have been concluded to be insensitive to amino acid signaling.^11^

**Figure 2.**
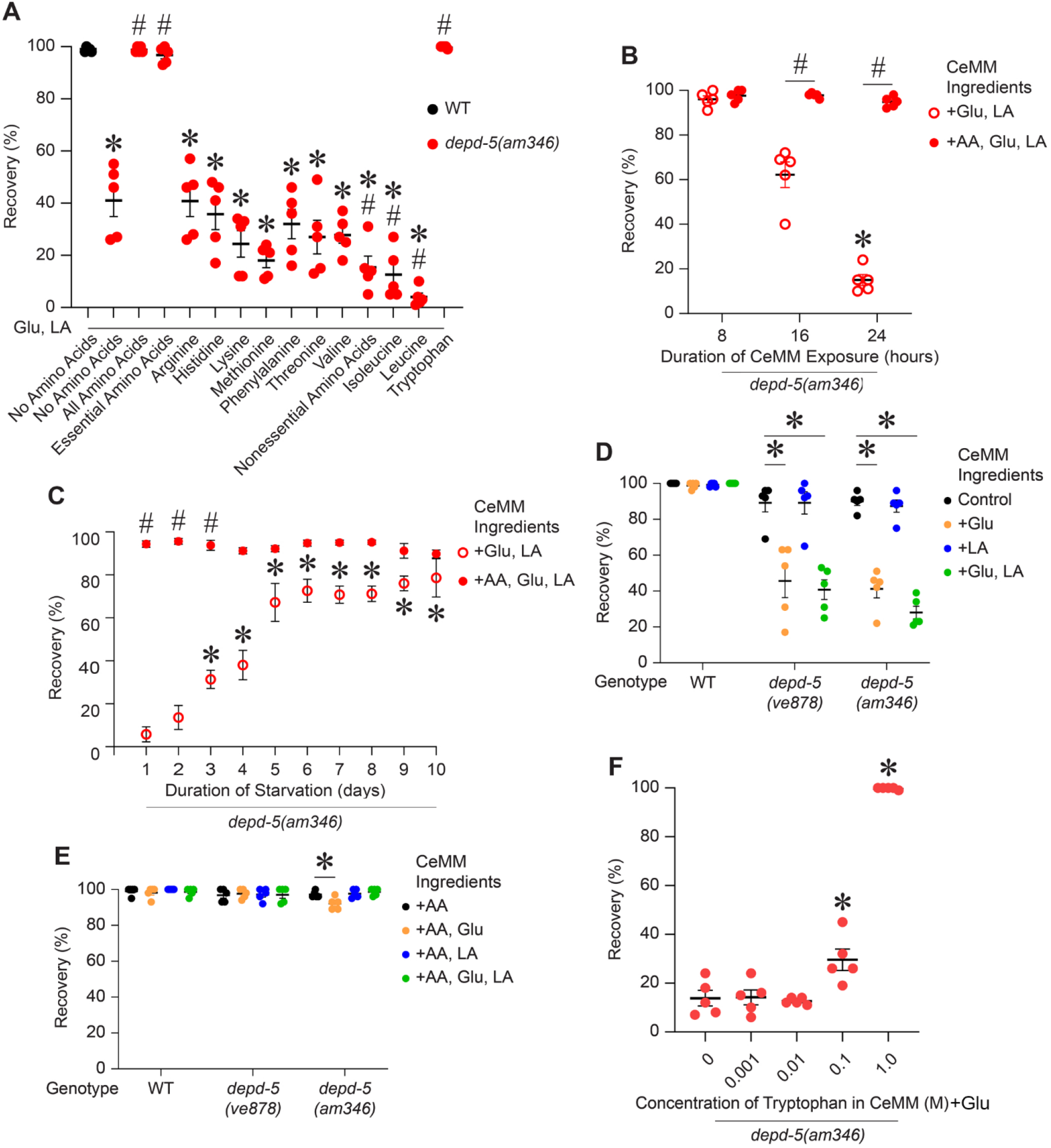
Glucose impairs starvation recovery and tryptophan restores recovery competence in *depd-5(lf)* worms. **(A)** L1 larvae were starved for 24 hours, exposed to CeMM minimal formulations supplemented with glucose (Glu), linoleic acid (LA) and various amino acids as indicated for 24 hours, cultured on *E. coli* OP50 layered agar dishes for 48 hours, and scored for survival (following the schematic depicted in Figure S4A). The horizontal axis shows the amino acids used, and the vertical axis shows recovery of wild-type and *depd-5(am346)* animals. Bars and whiskers represent mean ± SEM (five independent experiments). Statistical differences were determined by ordinary one-way ANOVA followed by Tukey’s multiple-comparisons test. \**P* < 0.05 compared with the wild-type no amino acid condition. #*P* < 0.05 for the reduced recovery compared with the *depd-5(lf)* with no amino acid added condition. **(B)** L1 larvae from *depd-5(am346)* mutant strain were starved for 24 hours, exposed to CeMM minimal formulations supplemented with glucose (Glu) and linoleic acid (LA) or glucose, linoleic acid and amino acids (AA) for 8, 16, 24 hours, cultured on *E. coli* OP50 layered agar dishes for 48 hours, and scored for survival. The horizontal axis shows the duration of CeMM exposure and the vertical axis shows recovery of *depd-5(am346)* animals. Bars and whiskers represent mean ± SEM (five independent experiments). Statistical significance among time points was determined by Kruskal-Wallis test followed by Dunn’s multiple-comparisons test. \**P* < 0.05 vs the 8 hour time point and #*P* < 0.05 versus Glu+LA group at the same time point. **(C)** L1 larvae from *depd-5(am346)* mutant strain were starved for 1-10 days, exposed to CeMM minimal formulations supplemented with glucose (Glu) and linoleic acid (LA) or glucose, linoleic acid, and amino acids (AA) for 24 hours, cultured on *E. coli* OP50 layered agar dishes for 48 hours, and scored for survival. The horizontal axis shows the duration of starvation, and the vertical axis shows recovery of *depd-5(am346)* animals. Bars and whiskers represent mean ± SEM (five independent experiments). Statistical significance among starvation duration was determined by ordinary one-way ANOVA followed by Dunnett’s multiple-comparisons test. *P < 0.05 versus 1 day time point, and *#P* < 0.05 for comparisons between medium lacking or containing amino acids (+AA) at the same time point. **(D)** L1 larvae of the indicated genotypes were starved for 24 hours, exposed to CeMM minimal formulations supplemented with glucose (Glu), linoleic acid (LA), or both chemicals for 24 hours, cultured on *E. coli* OP50 layered agar dishes for 48 hours, and scored for survival. The horizontal axis shows the genotype and the vertical axis shows recovery of animals. Bars and whiskers represent mean ± SEM (five independent experiments). Data were not normally distributed and therefore analyzed by one-way ANOVA followed by Tukey’s multiple-comparisons test comparing nutrient conditions within each genotype. \**P* < 0.05 versus control within each genotype. **(E)** L1 larvae of the indicated genotypes were starved for 24 hours, exposed to CeMM minimal formulations supplemented with amino acids (AA), glucose (Glu), linoleic acid (LA), or both chemicals for 24 hours, and scored for survival. The horizontal axis shows the genotype, and the vertical axis shows the percent of live animals. Bars and whiskers represent mean ± SEM (five independent experiments). Statistical significance was determined by two-way ANOVA followed by Tukey’s multiple-comparisons test comparing nutrient conditions within each genotype. \**P* < 0.05 versus control within the genotype. **(F)** L1 larvae from *depd-5(am346)* mutant strain were starved for 24 hours, exposed to CeMM minimal formulations supplemented with glucose (Glu) and various concentrations of tryptophan for 24 hours, cultured on *E. coli* OP50 layered agar dishes for 48 hours, and scored for survival. The horizontal axis shows the concentration of tryptophan, and the vertical axis shows recovery of *depd-5(am346)* animals. Bars and whiskers represent mean ± SEM (five independent experiments). Statistical significance was determined by ordinary one-way ANOVA followed by Dunnett’s multiple-comparisons test with each tryptophan concentration compared with the 0 M control. \**P* < 0.05.

To determine if a specific amino acid was sufficient to restore recovery competence, we tested a mixture of essential amino acids, a mixture of nonessential amino acids, and individual amino acids. A mixture of essential amino acids restored starvation recovery, whereas a mixture of nonessential amino acids did not (**Figure 2A**). Testing individual amino acids revealed that tryptophan, but no other single amino, was sufficient to restore recovery competence in *depd-5(am346)* worms (**Figure 2A**). Supplementation with isoleucine, leucine and a mixture of non-essential amino acids reduced survival compared with no amino acids (**Figure 2A**).

### A Time Course of Impaired Recovery After Starvation in *depd-5* Mutant Worms

To determine how rapidly failure of recovery competence sets in, we starved *depd-5(am346)* mutants for 24 hours and then incubated for 8, 16, or 24 hours in CeMM containing glucose and linoleic acid and analyzed survival after 48 hours of culture on agar plates overlaid with OP50 *E. coli*. Recovery competence was not affected after exposure to glucose and linoleic acid for 8 hours, was partially impaired after 16 hours, and was significantly reduced at 24 hours (**Figure 2B**). Addition of amino acids preserved recovery competence throughout the interval. Thus, glucose and linoleic acid cause a time dependent decrease in recovery competence over 24 hours.

To examine recovery competence as a function of starvation duration, we starved *depd-5(am346)* mutants for 1-10 days prior to recovery in CeMM minimal medium containing glucose and linoleic acid for 24 hours and analyzed survival after 48 hours of culture on agar plates overlaid with OP50 *E. coli*. *depd-5(lf)* mutants displayed minimal recovery after days 1 and 2. After 3 and 4 days the mutants displayed partial recovery, and after 5-10 days the mutants displayed substantial recovery (**Figure 2C**). This was a surprising observation, since prolonged starvation causes less recovery in wild-type animals compared with shorter durations of starvation,^9^ likely secondary to ongoing depletion of nutrients.^4^ Addition of amino acids maintained recovery competence throughout the day 1-10 interval (**Figure 2C**). Together, these findings demonstrate that sensitivity to glucose and linoleic acid refeeding is lost with prolonged starvation in *depd-5(am346)* mutant worms.

### Recovery Competence Depends on Metabolizable Energy Input

To evaluate the contributions of glucose and linoleic acid, we tested these nutrients separately and together. Recovery following starvation was impaired when *dedp-5(am346)* worms were exposed to glucose alone but not linoleic acid alone; provision of amino acids rescued this defect in recovery competence (**Figure 2D, E**). Similar results were observed with the *depd-5(ve878)* mutant (**Figure 2D, E**). In subsequent experiments, we studied *depd-5(am346)* strain which is annotated as d*epd-5(lf)* to indicate a loss-of-function allele. To analyze the dose dependence of tryptophan, we measured the effects of supplementation with 0.001, 0.01, 0.1 and 1.0 M tryptophan together with glucose in CeMM minimal after starvation and assessed recovery in d*epd-5(lf)* worms 48 hours later after plating on *E. coli*. The lowest two concentrations did not enhance recovery from starvation; 0.1 M caused a partial effect, whereas 1.0 M caused full recovery from starvation (**Figure 2F**). Thus, the effects of tryptophan are dose sensitive.

Glucose is the primary energy source and sole carbohydrate in CeMM.^12^ To focus on the mechanism of glucose toxicity, we tested additional metabolizable nutrients, including metabolites generated during glucose metabolism. Fructose, pyruvate, acetate, and ATP all significantly impaired recovery of *depd-5(lf)*, whereas tryptophan restored recovery competence when added to all these substrates (**Figure 3A-E**). These findings suggested that the impairment of starvation recovery was not specific to glucose, raising the possibility that recovery competence can be impaired by a variety of metabolizable nutrient inputs.

**Figure 3.**
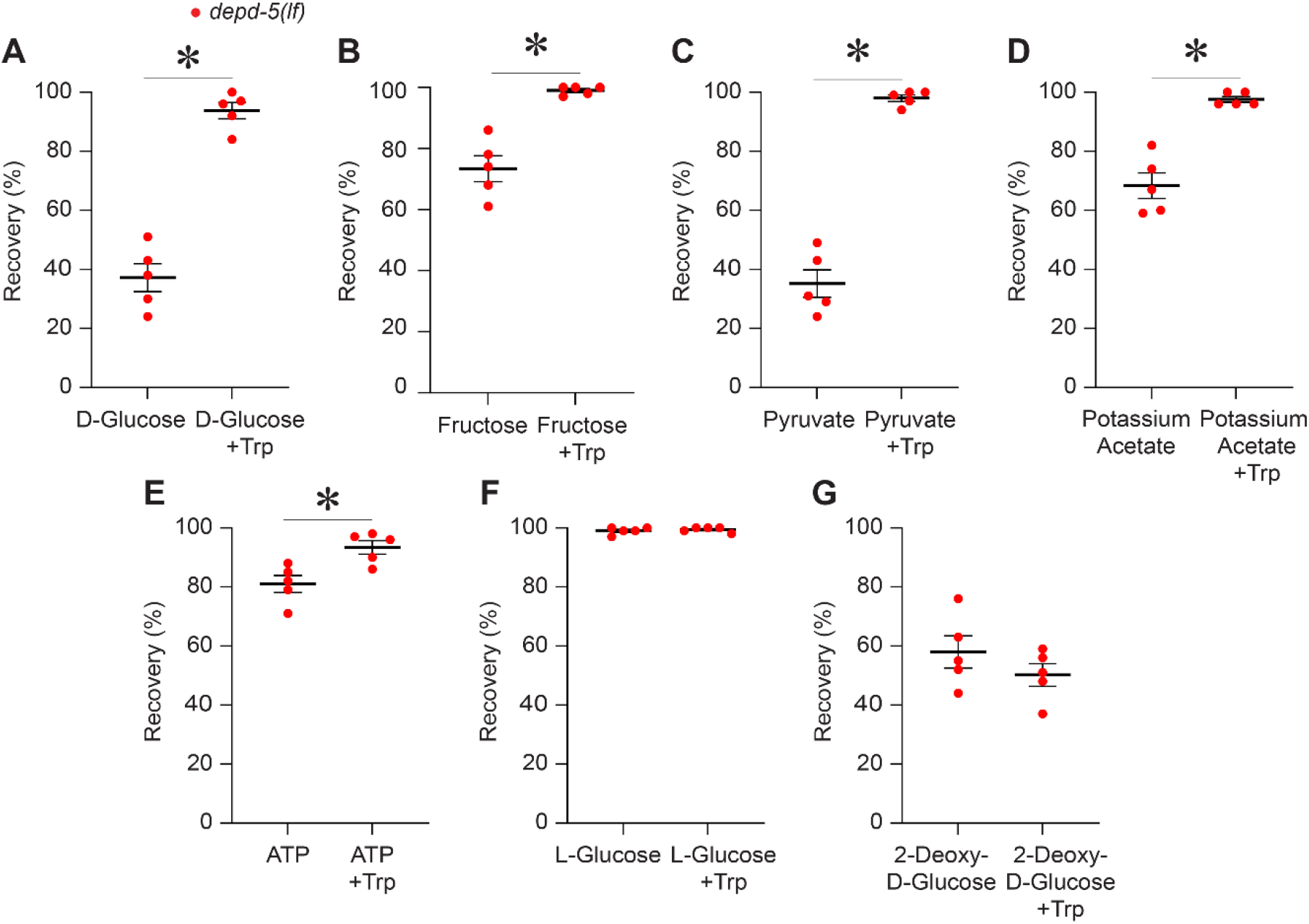
Metabolizable Energy in CeMM reduces Recovery Competence in *depd-5(lf)* worms. **(A-G)** L1 larvae from *depd-5(lf)* mutant strain were starved for 24 hours, exposed to CeMM minimal formulations supplemented with D-glucose (180 mM, A), fructose (180 mM, B), pyruvate (180 mM, C), potassium acetate (50 mM, D), ATP (10 mM, E), L-glucose (180 mM, F), or 2-deoxy-D-glucose (150 mM, G), with or without tryptophan (1 mM) for 24 hours, then cultured on *E. coli* OP50 layered agar dishes for 48 hours and scored for survival. The horizontal axis shows the chemicals added to CeMM and the vertical axis shows recovery of *depd-5(lf)* animals. Bars and whiskers represent mean ± SEM (five independent experiments). Statistical significance was determined using *t*-test; \**P* < 0.05.

We tested two glucose analogs with distinct metabolic properties. L-glucose, a stereoisomer that is not efficiently metabolized, did not have a significant effect (**Figure 3F**). To determine if productive glucose utilization was required, we supplemented with 2-deoxy-D-glucose, a glucose analog that inhibits productive glycolysis. This chemical did impair recovery in starved *depd-5(lf)* worms, but tryptophan did not rescue the effect (**Figure 3G**). Together, these findings suggest that impaired starvation recovery competence requires productive nutrient utilization rather than nutrient exposure alone.

### *depd-5* Genetically Interacts with *szt-2*, a Predicted Member of the KICSTOR Complex, and *raga-1*, a Predicted Regulator of mTORC1

Interestingly, the recovery of *depd-5(lf)* worms starved for 24 hours and exposed to CeMM minimal followed by 48 hours of culture on agar plates overlaid with OP50 *E. coli* appeared to modestly reduced (∼90%) as compared with WT (∼100%) even in the absence of glucose supplementation (**Figure 2D**). To rigorously evaluate this finding and examine genetic interactions in the pathway described for DEPDC-5 mediated signaling to mTOR, we performed additional studies. In mammals, the KICSTOR complex regulates GATOR1 localization and signaling (**Figure 4A**).^17^ Mammalian SZT2 is a component of the KICSTOR complex, and the *C. elegans* genome encodes a conserved *szt-2* gene. If the functions of the KICKSTOR Complex and GATOR1 are conserved in worms, we hypothesize that loss-of-function mutations of *szt-2* and *depd-5* will cause similar phenotypes (**Figure 4B**). To test this model, we obtained the *szt-2(gk554651)* allele with a predicted loss-of-function (*lf*). We starved *szt-2(lf)* mutants for 24 hours at the L1 larval stage and exposed these animals to CeMM minimal for 24 hours. The *szt-2(lf)* mutants displayed ∼25% lethality following treatment with CeMM minimal, which is not observed with *depd-5(lf)* mutants (**Figure S5A**). When these animals were transferred to agar plates with *E. coli* and scored for survival after 48 hours, *depd-5(lf)* worms demonstrate impaired recovery as compared with WT (∼100%; **Figure 4C**) both in the absence and presence of glucose. The *szt-2(lf)* mutants also displayed significantly impaired recovery under both conditions (**Figure 4C**), which was further impaired as compared with *depd-5(lf)* mutants. Tryptophan treatment significantly improved starvation recovery for *szt-2(lf)* mutants similar to *depd-5(lf)* mutants under both conditions. Thus, the *szt-2(lf)* mutant demonstrates a worse phenotype *depd-5(lf)* mutant in displaying partial lethality after CeMM treatment, but it is similar in that glucose further inhibits recovery from starvation and this effect is blunted by tryptophan.

**Figure 4.**
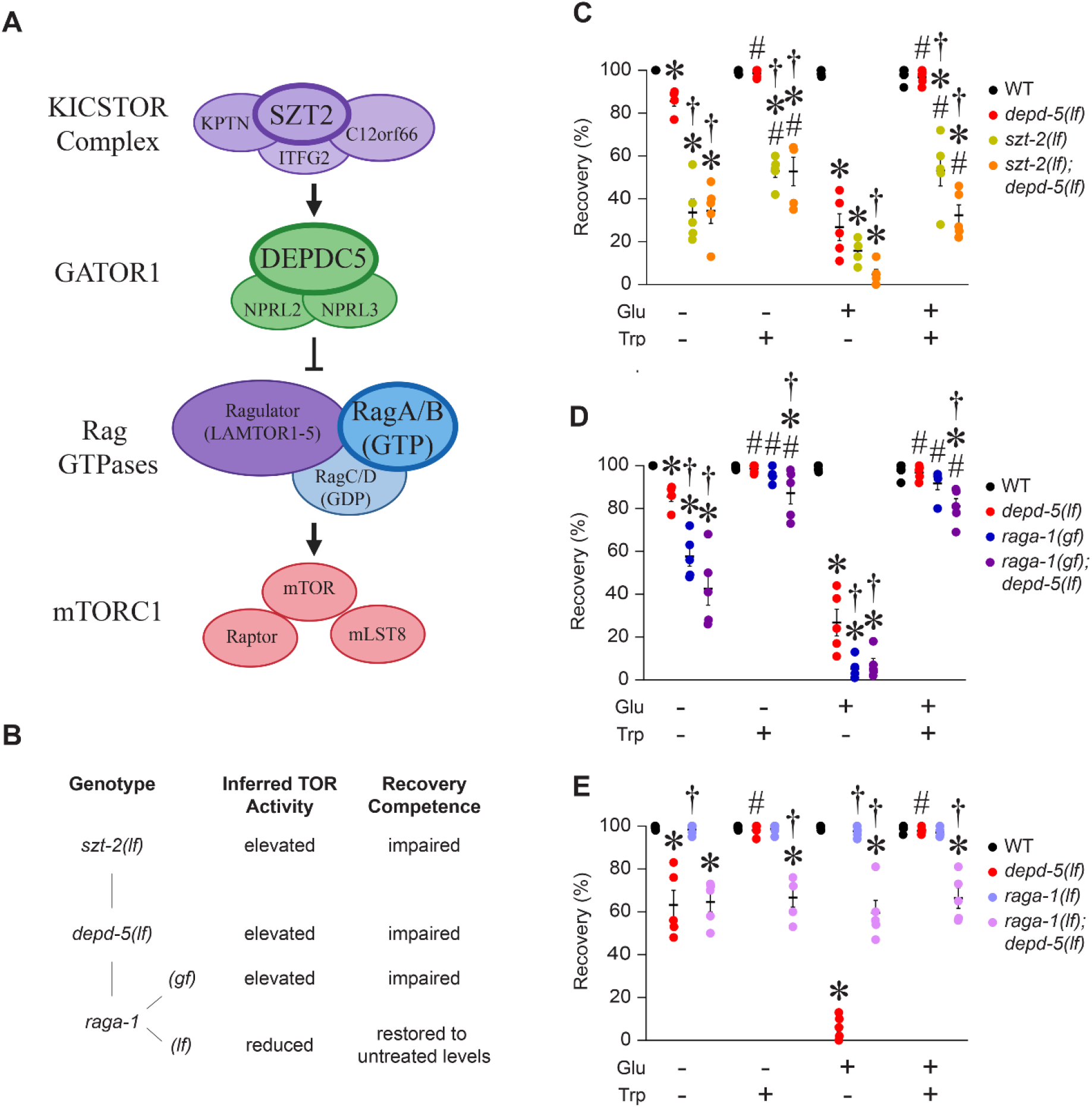
*depd-5* genetically interacts with *szt-2* and *raga-1*. **(A)** Schematic of the mammalian KICSTOR-GATOR1-Rag GTPase signaling pathway regulating mTORC1. KICSTOR is composed of four proteins, including SZT2 (purple); GATOR1 is composed of three proteins, including DEPDC5 (green); Rag GTPases is composed of multiple proteins, including RagA/B (blue); mTORC1 is composed of three proteins including mTOR. KICSTOR recruits GATOR1 to the lysosome where GATOR1 acts as a GTPase activating protein for RagA/B. Active Rag GTPases promote lysosomal recruitment of mTORC1. **(B)** Predicted recovery competence based on genotypes tested with inferred mTOR activity per schematic in panel A. **(C-E)** L1 larvae for indicated genotypes were starved for 24 hours, exposed to CeMM minimal formulations supplemented with glucose (Glu), tryptophan (Trp), or both chemicals for 24 hours, cultured on *E. coli* OP50 layered agar dishes for 48 hours and scored for survival. The horizontal axis shows the composition of CeMM and the vertical axis shows percent of animals that recover from starvation. Bars and whiskers represent mean ± SEM (five independent experiments). Percent recovery data were transformed using an arcsine square-root transformation before statistical analysis. Transformed values were analyzed by ordinary two-way ANOVA followed by Tukey’s multiple-comparisons test. \**P* < 0.05 vs. WT control within the respective nutrient group, †*P<0.05* vs. *depd-5(lf)* group within the same nutrient condition, and # indicates *P*<0.05 for Trp (+Tryptophan) vs. group without or with exposure to glucose (Glu) within the same genotype.

In mammals, DEPDC5 is part of the GATOR1 protein complex, which negatively regulates the activity of Rag GTPases (**Figure 4A**).^11^ If these functions are conserved in worms, we hypothesize that the *depd-5(lf)* mutation increases the activity of RagA and mTORC1 and thereby impairs starvation recovery in the presence of glucose (**Figure 4B**). This model predicts that a gain-of-function mutation of *raga-1* will phenocopy the *depd-5(lf)* phenotype, whereas a loss-of-function mutation of *raga-1* will suppress the *depd-5(lf)* mutant defects. To investigate this hypothesis, we obtained the *raga-1(viz128)* gain-of-function *(gf)* mutation, which increases RSKS-1 phosphorylation consistent with mTOR hyperactivation without affecting lifespan.^18^ We also obtained the *raga-1(ok386)* loss-of-function *(lf)* mutation, which extends lifespan and mTOR inhibition does not further extend lifespan in *raga-1(ok386)* animals,^19^ providing genetic evidence that RAGA-1 and mTOR act in a common pathway regulating longevity.

We starved *raga-1(gf)* mutants for 24 hours at the L1 larval stage and exposed these animals to CeMM minimal for 24 hours. *raga-1(gf)* mutants displayed little or no lethality following treatment with CeMM minimal, similar to *depd-5(lf)* mutants (**Figure S5B**). When these animals were transferred to standard medium with *E. coli* and scored for survival after 48 hours, the *raga-1(gf)* mutants also displayed significantly impaired recovery from starvation (∼10%) in the presence of glucose (**Figure 4D**). Furthermore, this defect was rescued by tryptophan (**Figure 4D**). Interestingly, the *raga-1(gf)* mutant also demonstrated impaired recovery similar to in *depd-5(lf)* and *szt-2(lf)* mutants under these experimental conditions in the absence of glucose supplementation, which was rescued by tryptophan treatment (**Figure 4C**).

These results are consistent with the model that *depd-5* is a negative regulator of *raga-1* in *C. elegans* and that the increase activity of *raga-1* is the cause of the failure to recover from starvation phenotype. To evaluate the interaction between the *raga-1(lf)* mutation and the *depd-5(lf)* mutation, we built the double mutant. We starved animals for 24 hours at the L1 larval stage and exposed these animals to CeMM minimal for 24 hours. *raga-1(lf);depd-5(lf)* double mutants displayed a low level of lethality following treatment with CeMM, which is not observed with single mutants (**Figure S5C**). These animals were transferred to standard medium with *E. coli* and scored for survival after 48 hours. In the presence of glucose, the *depd-5(lf)* single mutant displayed ∼10% starvation recovery. By contrast, the *raga-1(lf); depd-5(lf)* double mutant displayed a significantly higher level of ∼60% starvation recovery (**Figure 4C**). In this genetic background, tryptophan did not confer an additional survival advantage on the double mutant. Thus, reducing the activity of *raga-1* rescued the lethality of *depd-5(lf)* mutants, providing strong support for the model that the lethality in *depd-5(lf)* mutant animals is caused by activating RAGA-1.

### Tryptophan Availability Regulates Recovery Competence

In *C. elegans*, most tryptophan is metabolized by the enzyme tryptophan 2,3-dioxygenase (TDO-2) to generate kynurenine, while a smaller fraction is metabolized by the enzyme tryptophan hydroxylase (TPH-1) to generate 5-HTP and eventually serotonin^20^ (**Figure 5A**). The activity of these enzymes reduces the level of tryptophan and increases the level of metabolites. To investigate the basis for the tryptophan effects on starvation recovery, we administered exogenous L or D-tryptophan to the *depd-5(lf)* mutants. The biologically active L-tryptophan enantiomer restored recovery competence, whereas D-tryptophan did not (**Figure 5B**). Thus, restoration of recovery competence depends on stereospecific biological utilization of tryptophan.

**Figure 5.**
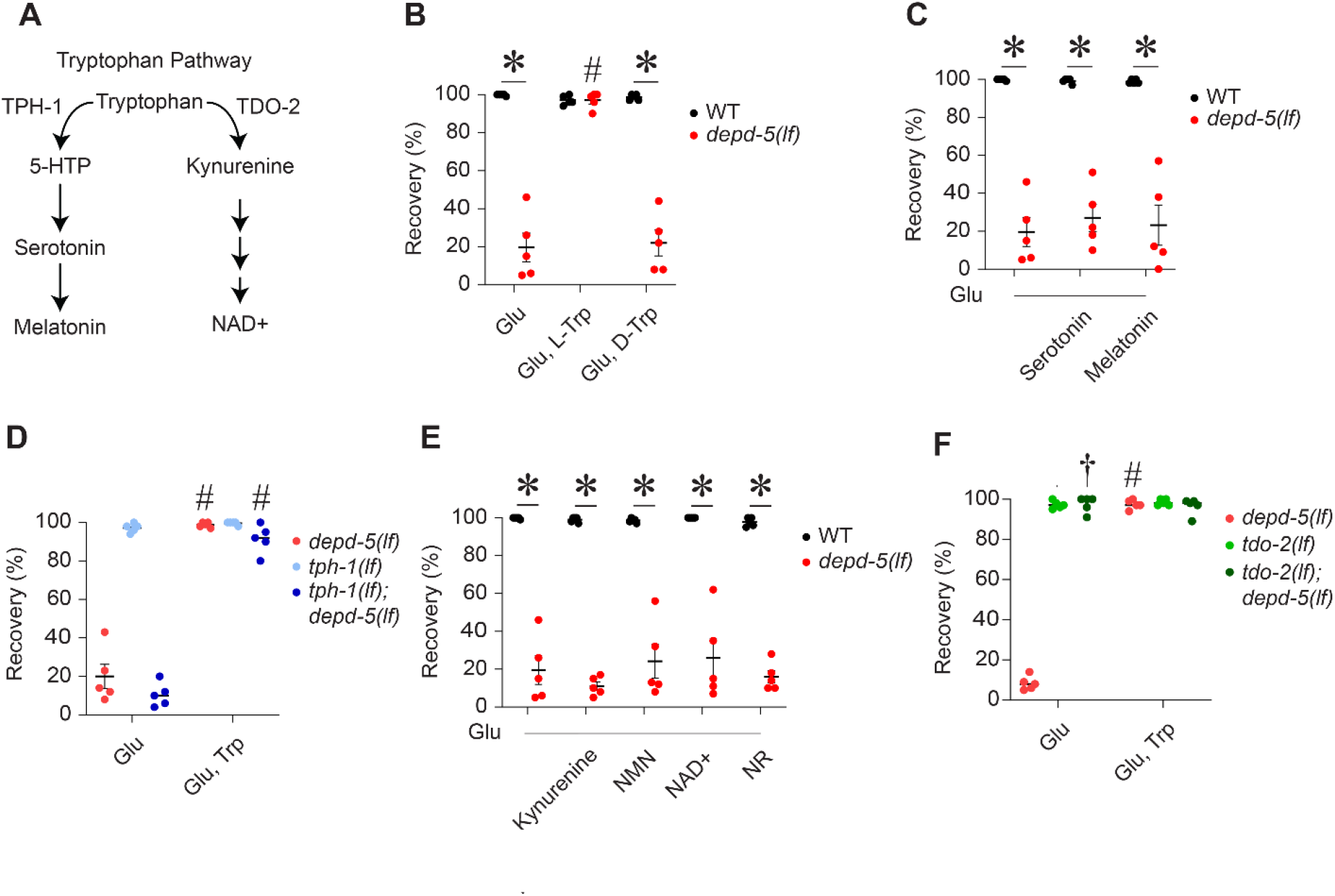
Tryptophan itself but not its metabolites regulate recovery competence. **(A)** Diagram depicting pathways of tryptophan metabolism through branches controlled by the enzymes TPH-1 and TDO-2. **(B)** L1 larvae of the indicated genotypes were starved for 24 hours, exposed to CeMM minimal formulations supplemented with glucose (Glu), glucose and L-tryptophan (Glu & L-Trp), glucose and D-tryptophan (Glu & D-Trp) for 24 hours, then cultured on *E. coli* OP50 layered agar dishes for 48 hours and scored for survival. All metabolites except glucose were added as 1 mM final concentration. The horizontal axis shows the chemicals added to CeMM minimal medium; the vertical axis shows recovery of wild-type and mutant animals, as indicated. Bars and whiskers represent the mean ± SEM (five independent experiments). Percentage recovery data were transformed using an arcsine square-root transformation before statistical analysis. Transformed values were analyzed by ordinary two-way ANOVA followed by Šídák’s multiple-comparisons test comparing genotypes within each nutritional condition. \**P* < 0.05. **(C)** L1 larvae of the indicated genotypes were starved for 24 hours, exposed to CeMM minimal formulations supplemented with glucose and serotonin, and glucose and melatonin for 24 hours, then cultured on *E. coli* OP50 layered agar dishes for 48 hours and scored for survival. All metabolites except glucose were added as 1 mM final concentration. The horizontal axis shows the chemicals added to CeMM minimal medium; the vertical axis shows recovery of wild-type and mutant animals, as indicated. Bars and whiskers represent the mean ± SEM (five independent experiments). Statistical significance was determined using two-tailed paired t-tests. **(D)** L1 larvae of the indicated genotypes were starved for 24 hours, exposed to CeMM minimal formulations supplemented with glucose with or without tryptophan for 24 hours, then cultured on *E. coli* OP50 layered agar dishes for 48 hours and scored for survival. Percentage recovery data were transformed using an arcsine square-root transformation before statistical analysis. Transformed values were analyzed by ordinary two-way ANOVA followed by Tukey’s multiple-comparisons test comparing genotypes within each nutrient condition. \**P* < 0.05 vs. wild type control. # *P* < 0.05 vs. glucose treated group for the same genotype. **(E)** L1 larvae of the indicated genotypes were starved for 24 hours, exposed to CeMM minimal formulations supplemented with glucose and kynurenine, glucose and nicotinamide mononucleotide (NMN), glucose and nicotinamide adenine dinucleotide (NAD+), glucose and nicotinamide riboside (NR) for 24 hours, then cultured on *E. coli* OP50 layered agar dishes for 48 hours and scored for survival. All metabolites except glucose were added as 1 mM final concentration. The horizontal axis shows the chemicals added to CeMM minimal medium; the vertical axis shows recovery of wild-type and mutant animals, as indicated. Bars and whiskers represent the mean ± SEM (five independent experiments). Statistical significance was determined using two-tailed paired t-tests. **(F)** L1 larvae of the indicated genotypes were starved for 24 hours, exposed to CeMM minimal formulations supplemented with glucose with or without tryptophan for 24 hours, then cultured on *E. coli* OP50 layered agar dishes for 48 hours and scored for survival. Percentage recovery data were transformed using an arcsine square-root transformation before statistical analysis. Transformed values were analyzed by ordinary two-way ANOVA followed by Šídák’s multiple-comparisons test comparing nutritional conditions within each genotype. †*P* < 0.05 vs. depd-5(lf). #*P* < 0.05 vs. glucose treated group for the same genotype.

If a downstream metabolite of tryptophan is responsible for the starvation recovery activity, then (1) direct administration of that chemical might have an effect like tryptophan and (2) a mutation that blocks formation of the metabolite should cause resistance to the beneficial effects of tryptophan in *depd-5(lf)* mutants. By contrast, if tryptophan itself is responsible for the starvation recovery activity, then (1) direct administration of chemical metabolites will not have an effect like tryptophan and (2) a mutation that blocks tryptophan metabolism might cause tryptophan to accumulate, thereby suppressing the *depd-5(lf)* mutant phenotype. To test this model, we first focused on the *tph-1* branch of tryptophan metabolism. We supplemented CeMM minimal with serotonin or melatonin and tested survival in the starvation recovery assay; these chemicals did not increase recovery competence (**Figure 5C**). We starved *tph-1(lf)* single mutants and *tph-1(lf); depd-5(lf)* double mutants for 24 hours at the L1 larval stage and exposed these animals to CeMM minimal for 24 hours. The single and double mutants displayed no lethality following treatment with CeMM minimal (**Figure S6A**). When these mutant animals were transferred to standard medium with *E. coli* and scored for survival after 48 hours, the *tph-1(lf); depd-5(lf)* double mutants displayed significantly impaired recovery from starvation (∼10%) in the presence of glucose, mimicking the *depd-5(lf)* worms (**Figure 5D**). Furthermore, this defect was rescued by tryptophan (**Figure 5E**). Thus, metabolites of tryptophan generated by the activity of TPH-1 do not appear to mediate the starvation recovery activity of tryptophan.

To evaluate the *tdo-2* branch of tryptophan metabolism, we supplemented CeMM minimal with kynurenine, the NAD+ precursors nicotinamide mononucleotide (NMN) or nicotinamide ribose (NR), and NAD+. None of these chemicals increased recovery competence, suggesting that metabolites of tryptophan generated by the activity of TDO-2 do not mediate the starvation recovery activity of tryptophan (**Figure 5E**). To examine genetic interactions, we starved *tdo-2(lf)* single mutants and *depd-5(lf); tdo-2(lf)* double mutants for 24 hours at the L1 larval stage and exposed these animals to CeMM minimal for 24 hours. The single and double mutants displayed little or no lethality following treatment with CeMM minimal (**Figure S6B**). When these mutant animals were transferred to standard medium with *E. coli* and scored for survival after 48 hours, the *depd-5(lf); tdo-2(lf)* double mutants displayed full recovery from starvation in the presence of glucose (**Figure 5F**). Thus, the *tdo-2* loss-of-function mutation suppresses the *depd-5(lf)* phenotype.

The striking interaction between *tdo-2* and *depd-5* suggests that the *tdo-2* mutation causes the accumulation of tryptophan, which in turn rescues the *depd-5(lf)* defect. To test this model, we measured tryptophan levels in worm extracts by targeted metabolomics with mass spectrometry. *depd-5* mutants treated with glucose displayed a tryptophan level of ∼3.2; treatment with glucose and tryptophan increased this level 20-fold to ∼61. The *depd-5(lf) tdo-2(lf)* double mutant treated with glucose displayed a tryptophan level of ∼74, similar to the level achieved by supplementation of tryptophan (**Table 1**). These findings demonstrate that the *tdo-2(lf)* mutation causes a dramatic increase in the level of tryptophan. To rule out the possibility that production of kynurenine by tryptophan breakdown via TDO-2 contributes to loss of recovery competence in *depd-5(lf)* worms, we starved *depd-5(lf) tdo-2(lf)* double mutant worms for 24 hours to CeMM minimal at the L1 larval stage and exposed these animals to CeMM minimal for 24 hours supplemented with increasing concentrations of kynurenine. There was no lethality following treatment with kynurenine in CeMM minimal (**Figure S6C**) or 48 hours after transfer to standard medium with *E. coli* (**Figure S6D**). These findings indicate that increased tryptophan levels are the likely reason that loss of TDO-2 activity restores recovery competence in the *depd-5(lf)* defect.

**Table 1.**
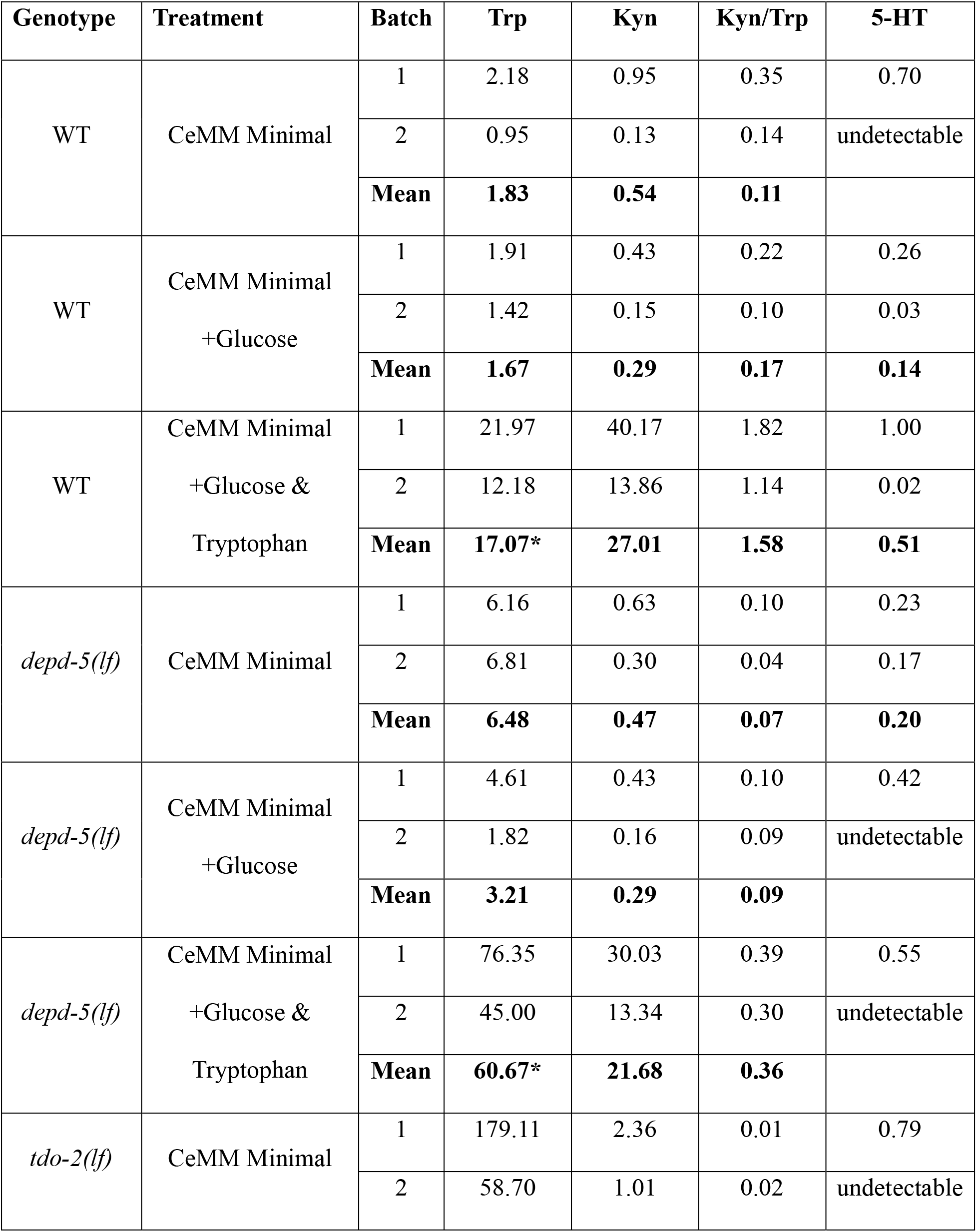
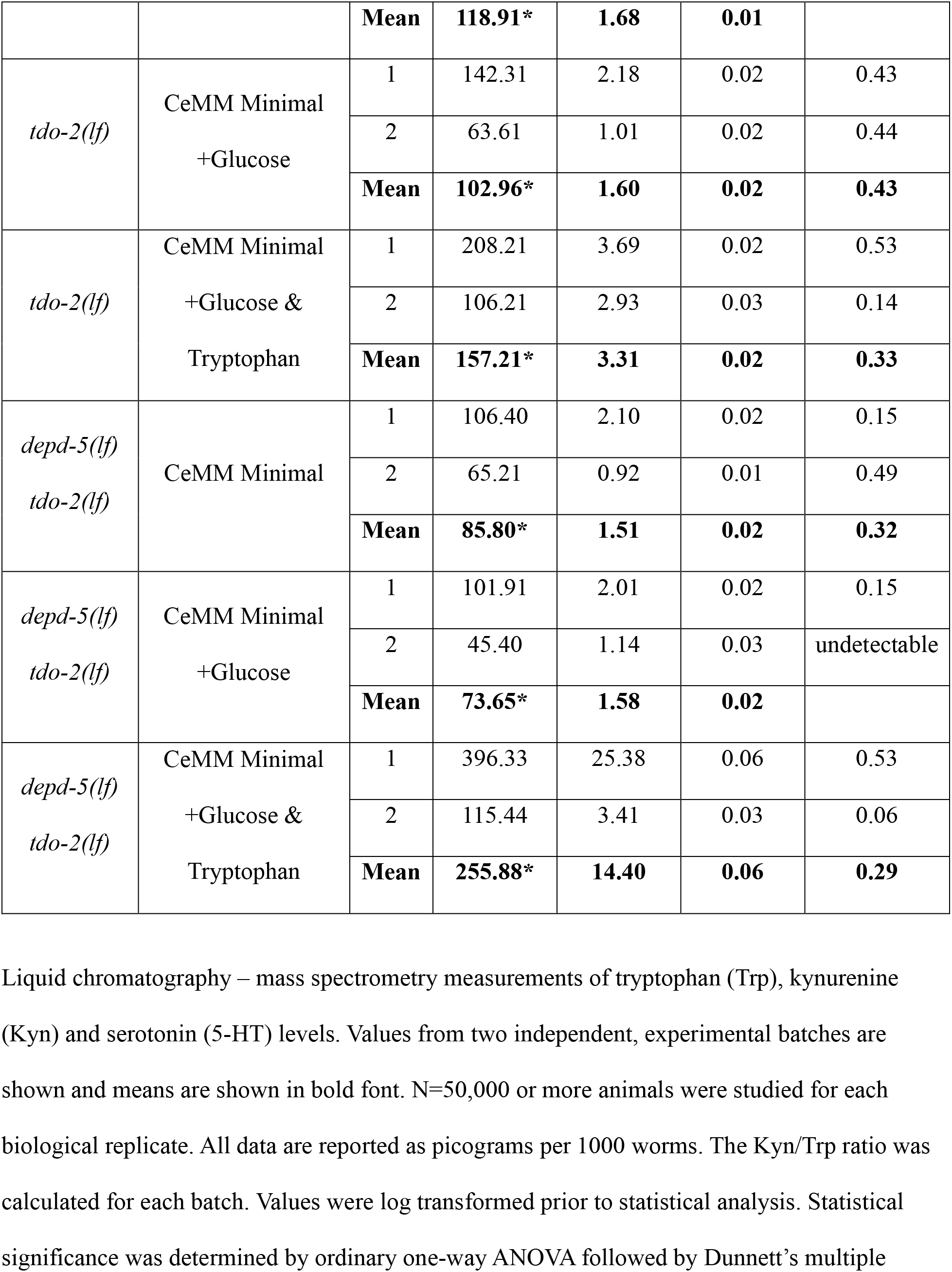
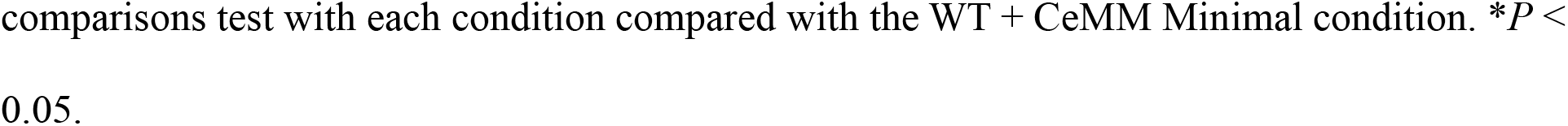
Mass spectrometry analysis of tryptophan metabolites with exposure to various CeMM formulations for 24 hours following 24 hours of starvation.

| **Genotype** | **Treatment** | **Batch** | **Trp** | **Kyn** | **Kyn/Trp** | **5-HT** |
| --- | --- | --- | --- | --- | --- | --- |
| WT | CeMM Minimal | 1 | 2.18 | 0.95 | 0.35 | 0.70 |
|  |  | 2 | 0.95 | 0.13 | 0.14 | undetectable |
|  |  | **Mean** | **1.83** | **0.54** | **0.11** |  |
| WT | CeMM Minimal<br>+Glucose | 1 | 1.91 | 0.43 | 0.22 | 0.26 |
|  |  | 2 | 1.42 | 0.15 | 0.10 | 0.03 |
|  |  | **Mean** | **1.67** | **0.29** | **0.17** | **0.14** |
| WT | CeMM Minimal<br>+Glucose &<br>Tryptophan | 1 | 21.97 | 40.17 | 1.82 | 1.00 |
|  |  | 2 | 12.18 | 13.86 | 1.14 | 0.02 |
|  |  | **Mean** | **17.07\*** | **27.01** | **1.58** | **0.51** |
| *depd-5(lf)* | CeMM Minimal | 1 | 6.16 | 0.63 | 0.10 | 0.23 |
|  |  | 2 | 6.81 | 0.30 | 0.04 | 0.17 |
|  |  | **Mean** | **6.48** | **0.47** | **0.07** | **0.20** |
| *depd-5(lf)* | CeMM Minimal<br>+Glucose | 1 | 4.61 | 0.43 | 0.10 | 0.42 |
|  |  | 2 | 1.82 | 0.16 | 0.09 | undetectable |
|  |  | **Mean** | **3.21** | **0.29** | **0.09** |  |
| *depd-5(lf)* | CeMM Minimal<br>+Glucose &<br>Tryptophan | 1 | 76.35 | 30.03 | 0.39 | 0.55 |
|  |  | 2 | 45.00 | 13.34 | 0.30 | undetectable |
|  |  | **Mean** | **60.67\*** | **21.68** | **0.36** |  |
| *tdo-2(lf)* | CeMM Minimal | 1 | 179.11 | 2.36 | 0.01 | 0.79 |
|  |  | 2 | 58.70 | 1.01 | 0.02 | undetectable |
|  |  | **Mean** | **118.91\*** | **1.68** | **0.01** |  |
| *tdo-2(lf)* | CeMM Minimal<br>+Glucose | 1 | 142.31 | 2.18 | 0.02 | 0.43 |
|  |  | 2 | 63.61 | 1.01 | 0.02 | 0.44 |
|  |  | **Mean** | **102.96\*** | **1.60** | **0.02** | **0.43** |
| *tdo-2(lf)* | CeMM Minimal<br>+Glucose &<br>Tryptophan | 1 | 208.21 | 3.69 | 0.02 | 0.53 |
|  |  | 2 | 106.21 | 2.93 | 0.03 | 0.14 |
|  |  | **Mean** | **157.21\*** | **3.31** | **0.02** | **0.33** |
| *depd-5(lf)*<br>*tdo-2(lf)* | CeMM Minimal | 1 | 106.40 | 2.10 | 0.02 | 0.15 |
|  |  | 2 | 65.21 | 0.92 | 0.01 | 0.49 |
|  |  | **Mean** | **85.80\*** | **1.51** | **0.02** | **0.32** |
| *depd-5(lf)*<br>*tdo-2(lf)* | CeMM Minimal<br>+Glucose | 1 | 101.91 | 2.01 | 0.02 | 0.15 |
|  |  | 2 | 45.40 | 1.14 | 0.03 | undetectable |
|  |  | **Mean** | **73.65\*** | **1.58** | **0.02** |  |
| *depd-5(lf)*<br>*tdo-2(lf)* | CeMM Minimal<br>+Glucose &<br>Tryptophan | 1 | 396.33 | 25.38 | 0.06 | 0.53 |
|  |  | 2 | 115.44 | 3.41 | 0.03 | 0.06 |
|  |  | **Mean** | **255.88\*** | **14.40** | **0.06** | **0.29** |

Liquid chromatography – mass spectrometry measurements of tryptophan (Trp), kynurenine (Kyn) and serotonin (5-HT) levels. Values from two independent, experimental batches are shown and means are shown in bold font. N=50,000 or more animals were studied for each biological replicate. All data are reported as picograms per 1000 worms. The Kyn/Trp ratio was calculated for each batch. Values were log transformed prior to statistical analysis. Statistical significance was determined by ordinary one-way ANOVA followed by Dunnett's multiple
comparisons test with each condition compared with the WT + CeMM Minimal condition. $*P < 0.05$ .

We described above the surprising observation that prolonged starvation in *depd-5(lf)* mutants improved recovery competence (**Figure 2C**). To test the model that this time-dependent improvement is due to the accumulation of tryptophan, we measured tryptophan levels after 7 days of starvation. *depd-5(lf)* worms demonstrated significantly elevated tryptophan levels on day 7 compared with wild type controls starved for 7 days with or without glucose supplementation (**Table 2**). These findings indicate that tryptophan itself, rather than downstream metabolites, causes increased recovery competence of *depd-5(lf)* mutants.

**Table 2.** Mass Spectrometry analysis of tryptophan metabolites with exposure to various CeMM formulations for 24 hours following 7 days of starvation.

| **Genotype** | **Treatment** | **Batch** | **Trp** | **Kyn** | **Kyn/Trp** | **5-HT** |
| --- | --- | --- | --- | --- | --- | --- |
| WT | CeMM Minimal | 1 | 3.04 | 0.22 | 0.07 | 0.26 |
|  |  | 2 | 2.55 | 0.17 | 0.07 | 0.37 |
|  |  | **Mean** | **2.79** | **0.19** | **0.07** | **0.31** |
| WT | CeMM Minimal<br>+Glucose | 1 | 1.15 | 0.10 | 0.09 | 0.49 |
|  |  | 2 | 1.13 | 0.10 | 0.09 | 0.47 |
|  |  | **Mean** | **1.14** | **0.10** | **0.09** | **0.48** |
| *depd-5(lf)* | CeMM Minimal | 1 | 26.68 | 1.48 | 0.06 | 0.37 |
|  |  | 2 | 32.23 | 1.93 | 0.06 | 0.56 |
|  |  | **Mean** | **29.46\*** | **1.70** | **0.06** | **0.47** |
| *depd-5(lf)* | CeMM Minimal<br>+Glucose | 1 | 18.91 | 1.22 | 0.07 | 0.80 |
|  |  | 2 | 20.28 | 1.00 | 0.05 | 0.70 |
|  |  | **Mean** | **19.60\*** | **1.11** | **0.06** | **0.75** |

Liquid chromatography – mass spectrometry measurements of tryptophan (Trp), kynurenine (Kyn) and serotonin (5-HT) levels. N=50,000 or more animals were studied for each biological replicate. All data are reported as picograms per 1000 worms. Values from two independent, experimental batches of worms starved for 7 days, their means are shown in bold font. The Kyn/Trp ratio was calculated for each batch. Statistical significance was determined by ordinary one-way ANOVA followed by Dunnett’s multiple-comparisons test with each condition compared with the WT CeMM Minimal condition. \* $P < 0.05$ .

### Nutrient and Stress-Sensing Pathways Differentially Regulate Recovery Competence

Our results indicate that availability of tryptophan, an amino acid, was strongly associated with recovery competence following starvation in the absence of *depd-5*, the worm ortholog of DEPDC5 which is a core component of canonical amino acid sensing complex (GATOR1).^11^ Accordingly, we asked if an alternate amino acid sensor such as GCN-2^21,22^ was required for the response to tryptophan. We obtained two independently derived *gcn-2(lf)* mutants^21^ and generated gcn-2*(lf); depd-5(lf)* double mutant worms. To examine genetic interactions, we starved *gcn-2(lf)* single mutants and gcn-2*(lf); depd-5(lf)* double mutants for 24 hours at the L1 larval stage and exposed these animals to CeMM minimal for 24 hours. The single and double mutants displayed little or no lethality following treatment with CeMM minimal supplemented with glucose or tryptophan (**Figure S7A**). When these mutant animals were transferred to standard medium with *E. coli* and scored for survival after 48 hours, the *gcn-2(lf); depd-5(lf)* double mutants displayed significantly impaired recovery from starvation in the presence of glucose, mimicking the *depd-5(lf)* worms (**Figure 6A**). Furthermore, this defect was rescued by tryptophan (**Figure 6A**). These data indicate that GCN-2 was not required for tryptophan-dependent restoration of recovery competence.

**Figure 6.**
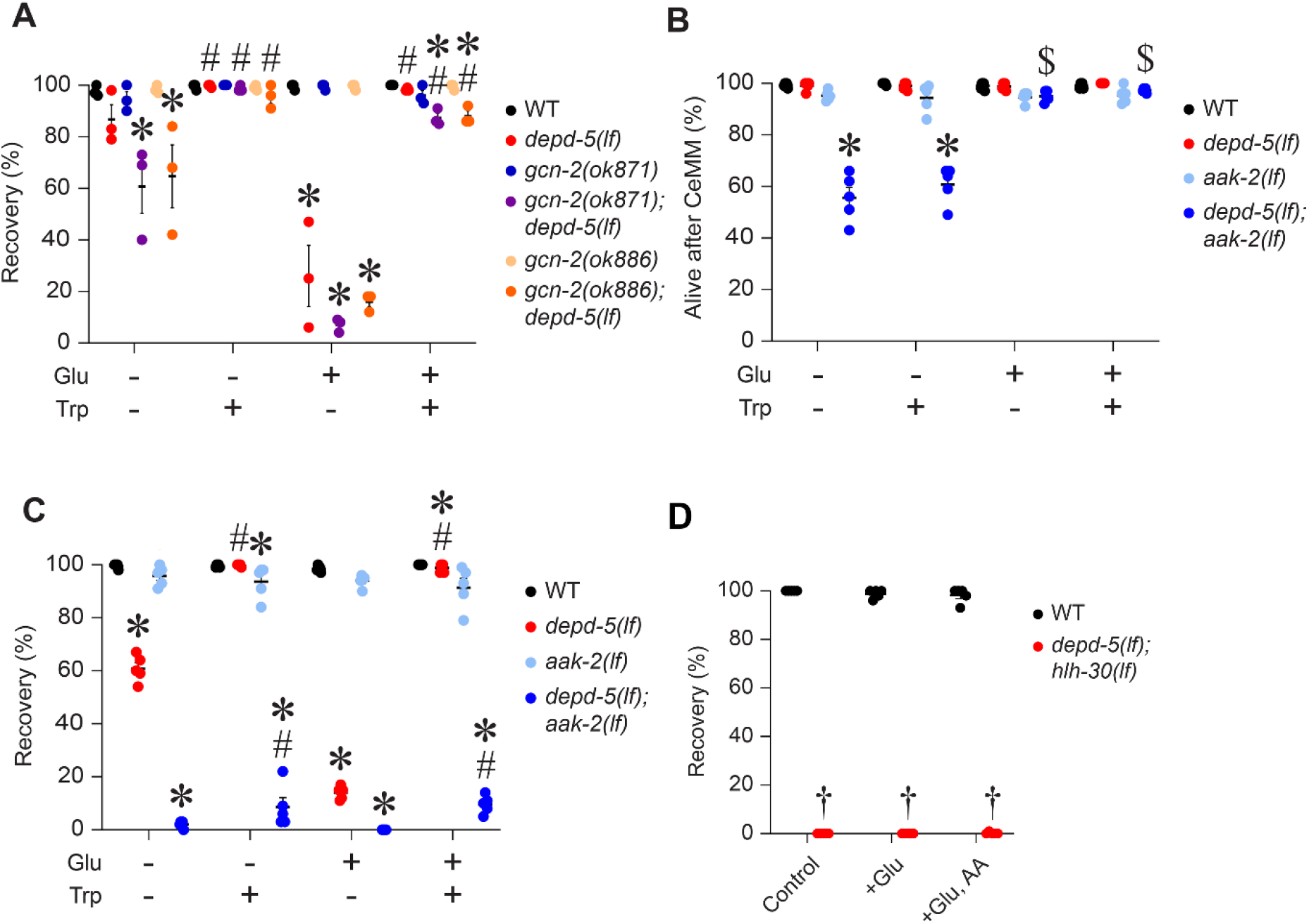
Amino acid and stress-sensing pathways differentially regulate recovery competence. **(A)** L1 larvae for indicated genotypes with *gcn-2(lf)* alone or in combination with *depd-5(lf)* were starved for 24 hours, exposed to CeMM minimal formulations supplemented with glucose (Glu), tryptophan (Trp), or both chemicals for 24 hours, then cultured on *E. coli* OP50 layered agar dishes for 48 hours, and scored for survival. The horizontal axis shows the composition of CeMM and the vertical axis shows percent of animals that recover from starvation. Bars and whiskers represent mean ± SEM (five independent experiments). The percent recovery data were transformed using an arcsine square-root transformation before statistical analysis. Transformed values were analyzed by ordinary two-way ANOVA followed by Tukey’s multiple-comparisons test comparing genotypes within each nutrient condition. \**P* < 0.05 versus wild type control and #*P*<0.05 for tryptophan group vs. same glucose treatment condition (without or with glucose) within the same genotype. **(B, C)** L1 larvae of worm strains with the indicated genotypes with *aak-2(lf)* and and in combination with *depd-5(lf),* and wild type controls were starved for 24 hours, exposed to CeMM minimal formulations supplemented with glucose (Glu) or glucose and tryptophan (Glu, Trp) for 24 hours, and immediately scored for survival (in B, assessed as Alive after CeMM (%)). These worms were then cultured on *E. coli* OP50 layered agar dishes for 48 hours and scored for survival (in C, assessed as Recovery (%)). The horizontal axis shows the composition of CeMM minimal and the vertical axis shows percent of animals that are alive after the CeMM treatment (B) or after being cultured on *E. coli* (C). Symbols represent individual experiments and the horizontal bars represent mean ± SEM (five independent experiments). Percentage survival data were transformed using an arcsine square-root transformation before statistical analysis. Transformed values were analyzed by ordinary two-way ANOVA followed by Tukey’s multiple-comparisons test comparing genotypes within each nutritional condition. \**P*<0.05 versus wild type within the same nutrient treated group; $*P*<0.05 for glucose treated group as compared with group without glucose treatment, either in the presence or absence of tryptophan; #*P*<0.05 for tryptophan group vs. same glucose treatment condition (without or with glucose) within the same genotype. **(D)** Recovery of wild-type and *depd-5(am346); hlh-30(lf)* animals following incubation in CeMM formulations containing glucose, glucose and amino acids or neither. Data are indicated as Mean ± SEM (from five independent experiments). † indicates 0% recovery in all independent experiments, and statistical analysis was not performed.

Given our observations indicating that recovery competence following starvation in the absence of *depd-5* is further impaired with provision of energy generating nutrients and ATP (**Figure 3A-E**), we asked if pathways that sense the cellular energic state influenced recovery. The conserved AAK-2/AMPK pathway responds to cellular energy availability and promotes adaptation to energetic stress.^23^ We asked whether loss of *aak-2* affected recovery competence in *depd-5(lf)* mutants. Accordingly, we generated *depd-5(lf);* aak-2*(lf)* double mutants and starved them along with *aak-2(lf)* single mutants for 24 hours at the L1 larval stage and exposed to CeMM minimal for 24 hours. Interestingly, the *depd-5(lf);* aak-2*(lf)* double mutant displayed reduced survival as compared with wild type while survival of the *aak-2(lf)* and *depd-5(lf)* single mutants was not impaired (**Figure 6B**). Importantly, the lethality in *depd-5(lf);* aak-2*(lf)* double mutants at this stage was rescued by feeding glucose but not tryptophan (**Figure 6B**). This is consistent with prior observations where *aak-2(lf)* mutants demonstrate increased mortality with prolonged starvation during L1 arrest, and these worms could be rescued by providing glucose or ethanol as energy sources during this period.^24^ Taken together, loss of *aak-2* is synthetic lethal with loss of *depd-5* and confers impaired survival after starvation, which is rescued by glucose but not tryptophan. Thereafter, when these worms were transferred to *E. coli* layered agar plates and survival was assessed 48 hours later, the *depd-5(lf);* aak-2*(lf)* double mutants failed to recover (**Figure 6C**). This failure to recover was worse in the double mutant as compared with the *depd-5(lf)* single mutants in the absence of glucose, and similar to the *depd-5(lf)* single mutants in the presence of glucose (**Figure 6C**). Remarkably, tryptophan only had a modest effect on survival in the *depd-5(lf);* aak-2*(lf)* double mutants and failed to restore recovery in the double mutant to levels observed in the *depd-5(lf)* single mutant supplemented with tryptophan (**Figure 6C**). These findings establish that tryptophan fails to restore recovery competence in the absence of AAK-2 in worms lacking *depd-5* suggesting that tryptophan may confer beneficial effects on recovery competence via AMPK signaling (**Figure 6C**).

We next tested if HLH-30 was essential for recovery with tryptophan supplementation in starved *depd-5(lf)* mutants fed with glucose, by studying the *depd-5(lf); hlh-30(lf)* worms, which demonstrate impaired recovery competence with glucose and linoleic acid exposure following starvation (**Figure 1D**). We subjected *depd-5(lf); hlh-30(lf)* and WT worms to 24 hours starvation at the L1 larval stage and exposed to CeMM minimal with and without glucose for 24 hours. The *depd-5(lf); hlh-30(lf)* mutant worms did not demonstrate impaired survival at this stage (**Figure S7B**). However, supplementation with glucose was sufficient to impair recovery competence in *depd-5(lf); hlh-30(lf)* worms after 48 hours of refeeding with *E. coli* on agar plates (**Figure 6D**). Importantly, provision of amino acid cocktail which included 1mM tryptophan was unable to rescue these worms (**Figure 6D**). Taken together with our previously published observations,^9^ these data indicate that HLH-30 is required for recovery competence after starvation and indicate the tryptophan mediated rescue requires intact HLH-30 signaling.

### DEPDC5 Regulates mTOR Activity During Starvation in Human Cells

Our genetic results suggest a role for *depd-5* in restraining mTOR activation via *raga-1* inhibition upon starvation in worms, as shown in **Figure 4A, B**. To test whether introducing a mutation at an analogous location in the human *DEPDC5* ortholog will also affect mTOR activation, we modeled this genetic lesion using CRISPR targeting. This mutation resulted in loss of DEPDC5 protein in HeLa cells (**Figure S8A**) generating a DEPDC5 knockout HeLa cell line. We compared the state of mTOR activation in wild-type and DEPDC5 mutant cells during starvation by measuring phosphorylation of the mTOR, and its targets, S6K and 4EBP-1. Starvation rapidly reduced S6K phosphorylation in wild-type cells indicating rapid mTOR inactivation **(Figure 7A, B).** In contrast, loss of DEPDC5 delayed the decline S6K phosphorylation with starvation in DEPDC5 mutant cells. Phosphorylation of 4EBP1 also showed a slower rate of decline in DEPDC5 mutant as compared with WT cells, while phosphorylation of mTOR did not demonstrate a similar trend without statistically significant differences during the same interval (**Figure 7A, S8B, C**).

**Figure 7.**
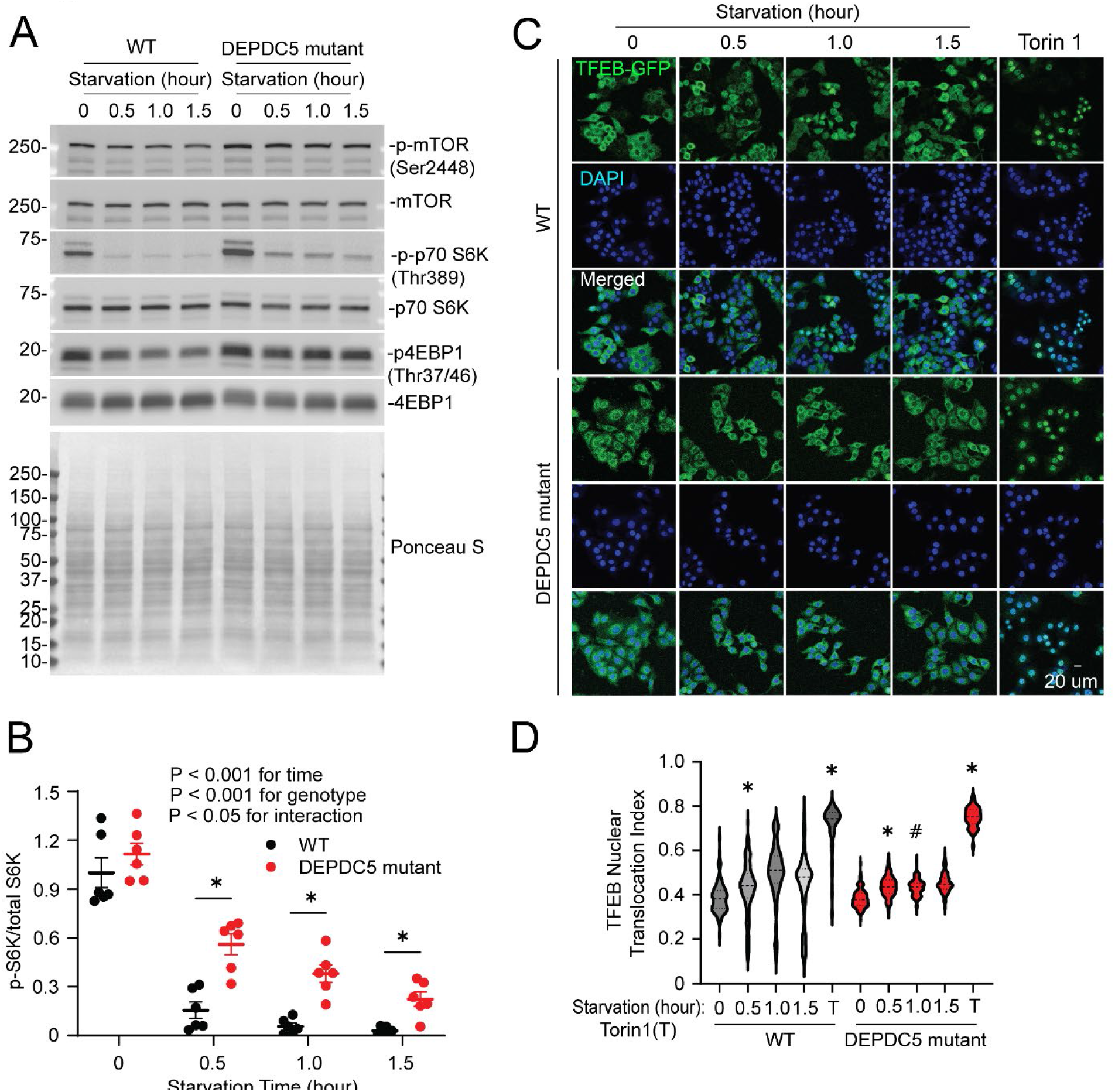
DEPDC5 restrains mTOR activity during starvation in human cells. **(A)** Representative immunoblots of phosphorylated and total mTOR, p70 S6K and 4EBP1 in wild-type and DEPDC5 mutant HeLa cells following starvation for the indicated times. Ponceau S staining was performed to assess total protein abundance. (**B**) Quantification of S6K phosphorylation relative to total S6K in wild-type and DEPDC5 mutant HeLa cells following starvation for the indicated times (x-axis). Data are from six independent experiments and expressed as fold over WT samples at T=0 hours. Statistical significance was determined by ordinary two-way ANOVA followed by Tukey’s multiple comparisons test. \**P* < 0.05. (**C**) Representative images of TFEB-GFP localization in wild-type and DEPDC5 knockout HeLa cells under fed conditions, following starvation for the indicated times, or following treatment with 500 nM Torin 1. DAPI staining identifies nuclei. (**D**) Quantification of TFEB nuclear localization under the conditions shown in (C). Statistical significance was determined using a Kruskal-Wallis test followed by Dunn’s multiple-comparisons test. N=100-180 nuclei were assessed per condition. \**P* < 0.05 compared with the fed condition of the same genotype and #*P*<0.05 vs. the identical time point in wild type cells.

The delayed suppression of S6K and 4EBP1 phosphorylation upon starvation in DEPDC5 mutant cells suggested that loss of DEPDC5 leads to unrestrained mTOR activity with starvation. HLH-30/TFEB plays a central role in the starvation response in *C. elegans*^9^ and mammalian TFEB is regulated by mTOR during starvation.^25^ We therefore examined if loss of DEPDC5 alters TFEB nuclear localization during starvation. TFEB-GFP subcellular localization was measured in wild-type and DEPDC5 mutant HeLa cells that were starved for 0.5, 1 and 1.5 hours and compared with Torin-treated cells as a positive control. Starvation increased nuclear localization of TFEB in wild-type cells, which was impaired in DEPDC5 mutant cells (**Figure 7D, E**). Torin treatment increased TFEB nuclear localization in both wild-type and DEPDC5 mutant cells indicating that TFEB is responsive to mTOR inhibition to undergo nuclear localization in the mutant, but its regulation under starvation conditions is altered by loss of DEPDC5. Together, these results support a conserved role for DEPD-5/DEPDC5 in regulating mTOR activity during starvation in worms and humans.

## Discussion

Our data reveal two key novel findings: 1) *depd-5* is a determinant of recovery competence following starvation, and recovery competence is worsened by glucose and energy generating substrates in its absence. 2) Tryptophan is essential in conferring recovery competence following starvation in the setting of unrestrained TOR activation. We have advanced our prior observations that introduction of nutrients after starvation does not guarantee recovery,^9^ as glucose and other metabolizable forms of carbon exposed a loss of recovery competence in the absence of canonical amino acid sensing. Our data support a model in which *depd-5* functions upstream of *raga-1* to regulate recovery competence (**Figure 4B**). Loss of *depd-5* and loss of *szt-2* phenocopies the genetically increased RAGA activation, while loss of *raga-1* restored recovery competence in *depd-5(lf)* mutants during glucose refeeding. Furthermore, we have uncovered that increased tryptophan availability restores recovery competence in all these settings. Taken together, these observations also point to a unique role for tryptophan signaling that is independent of currently described amino acid sensing pathways.

We had previously established a *C. elegans* model of starvation and recovery where glucose and linoleic acid were specifically required for recovery competence in the starvation and recovery-sensitive *hlh-30(lf)* mutant.^9^ That nutrient-based rescue of an otherwise irrecoverable phenotype provided the experimental entry point for genetic investigation using an unbiased genetic screen because the mechanisms that come into play during the transition from starvation to recovery remain unknown.^26^ We identified *depd-5*, which encodes a subunit of GATOR1, for its requirement in the glucose and linoleic acid mediated rescue of *hlh-30(lf)*. Earlier genetic and biochemical studies asked how the nutrient state is communicated to TORC1.^11,27,28^ Our screen selected for the consequence of this transition going awry. When *depd-5(lf)* was removed from the *hlh-30(lf)* background, and tested under the conditions of the original screen, glucose reduced recovery competence. Thus, we independently identified this machinery as critical to coupling nutrient availability to cellular nutrient-state signaling. This convergence raises the possibility that the known role of *depd-5* in nutrient-state regulation extends beyond amino acid sensing and contributes to successful recovery through as yet undescribed pathways.

Another remarkable experimental observation that recovery competence is spontaneously restored with prolonged starvation in the absence of DEPD-5 (Figure 2C). The ability of organisms to recover from nutrient deprivation is a biologically meaningful outcome. Crucially, across very different species, post-starvation outcomes worsen with increasing starvation duration. Starvation can alter what happens after nutrients return, ranging from a delayed transition to recovery,^29–32^ impaired recovery despite viability.^32–34^ In the established examples, the duration of starvation progressively created or aggravated the conditions associated with compromised recovery. In contrast, our findings show that increased starvation duration progressively reduced the glucose-sensitive state that compromised recovery competence in the *depd-5* loss-of-function worms. A related paradox was observed in a very different experimental system caused by loss of SEACIT (GATOR1 ortholog) in yeast.^35^ There was a mismatch between perceived nutrient sufficiency and actual metabolic capacity, with sustained TORC1 activity during amino acid scarcity and a growth defect in glucose-replete medium. Similarly, loss of the SEACIT components Npr2/Npr3 results in failure to adequately shut down TORC1 during amino acid starvation.^27^ These findings provide a metabolic precedent where nutrient signaling becomes maladaptive during nutrient insufficiency. We used gain and loss of function mutants in the GATOR1-Rag signaling pathway to test whether altering nutrient-state signaling affected recovery competence. We found that *raga-1(gf)* and *szt-2(lf)* mutants exhibited glucose-dependent reductions in recovery competence that were restored by tryptophan while recovery competence remained unaffected by glucose in the *raga-1(lf)* mutant. Consistent with a conserved role for GATOR1 in responding to nutrient state, loss of DEPDC5 in mammalian cells delayed the reduction in TORC1 signaling following nutrient withdrawal. Our findings suggest that when nutrients become available after starvation, the metabolic demands imposed by refeeding are mismatched, glucose, though beneficial in one context, can be profoundly detrimental when GATOR1 is disrupted. Prolonged starvation did not simply worsen recovery capacity; it changed the physiological state in which glucose was detrimental.

Pathogenic loss-of-function variants in the GATOR1 subunits DEPDC5, NPRL2 and NPRL3 cause focal epilepsy.^36^ Our data suggest that increasing tryptophan bioavailability may be a strategy to counter adverse consequences of unrestrained mTOR activation that is observed in epilepsy and aging-related diseases where mTOR hyperactivation is observed.^37^ Interestingly, loss of neuronal DEPDC5 in mice impaired the normal response to nutrient fluctuations during fasting,^38^ and acute fasting reduces brain mTORC1 activity and seizure susceptibility in a DEPDC5-dependent manner.^38^ Whereas the above mentioned study examined entry into fasting, our forward genetic approach revealed a *depd-5* dependent phenotype during exit from starvation. Both experimental systems uncovered abnormal responses when nutrient conditions change. Therefore, nutrient-state responsiveness may represent a broader biological function of DEPDC5/GATOR1. Extending this relationship beyond the worm is inherently complex because the phenotype reflects the combination of genotype, physiological state and nutritional environment. This raises the question of whether the postprandial metabolic response to tryptophan differs in DEPDC5 deficient mammals especially when glucose or other carbohydrate availability changes. We know considerably more about how organisms enter and maintain starvation states than about what determines their capacity to exit successfully when nutrients return. We have shown that recovery competence depends upon the interaction between the physiological state reached during starvation and the nutrients encountered during refeeding. This concept extends beyond *depd-5*, glucose and tryptophan which revealed one example of this interaction. The question of what determines whether an organism can successfully make the transition from starvation to recovery has only begun to be tested.

## Methods

### *C. elegans* Strains and Maintenance

*C. elegans* hermaphrodites were cultured under standard laboratory conditions^39^ on nematode growth medium (NGM) agar dishes and fed the *Escherichia coli* strain *OP50* at 20°C. A complete list of strains used in this study is provided in **Table S4**.

### Preparation of Worm Genomic DNA

Lysis buffer was prepared using 50 μL of 10× ThermoPol Reaction Buffer (New England BioLabs, #B9004S, Ipswich, MA, USA), 3.5 μL of 20 mg/mL Proteinase K solution from the QIAGEN DNeasy Blood & Tissue Kit (QIAGEN, #69506, Germantown, MD, USA) and 450 μL nuclease-free water. Approximately 20 worms were transferred from NGM dishes into 20 μL of lysis buffer. Samples were incubated at 60°C 60 min followed by 95°C for 15 min in a thermal cycler.

### Standard PCR, Cycling Conditions and Analysis of PCR Products

PCR reactions were performed in 25 μL volumes containing the following reagents at their final concentrations: Taq Plus 2× Master Mix (Molecular Designs, #D124R, Birmingham, AL, USA) at 1× concentration along with 0.4 μM of each primer (IDT, Coralville, IA, USA), 1 μL of DNA and nuclease-free water. PCR cycling conditions were: 95°C 5 minutes then 30 cycles of 94°C 30 seconds, 64°C 30 seconds, 72°C 60 seconds/1 kb of the amplicon, then 72°C 5 min. PCR products were separated on 1% agarose gels containing ethidium bromide and visualized under UV illumination.

Detection of the *depd-5(am346)* mutant was performed as follows.

*depd-5(am346)* F: ATACGCACTCACTTCGGATG
*depd-5(am346)* R: TGAGCCATCGTAGGCTTCTC

The *depd-5(am346)* F and R primers amplify a 1,256 bp amplicon in both wild-type and *depd-5(am346)* mutants. The cycling conditions were modified to include a 90 sec elongation time. The PCR product was purified using the Standard DNA Cleanup and Concentration protocol found in the Monarch Spin PCR & DNA Cleanup Kit (New England BioLabs, #T1130S, Ipswich, MA, USA). The purified PCR product was sent for Sanger Sequencing (Genewiz from Azenta Life Sciences, South Plainfield, NJ, USA) to determine the genotype at the *depd-5(am346)* locus.

### Generation of Mutant Strains

Construction of the raga-1(viz128) II; depd-5(am346) III double mutant

The *raga-1(viz128) II; depd-5(am346) III* double mutant was constructed with *depd-5(am346)* males and a *raga-1(viz128)* hermaphrodite on NGM dishes seeded with OP50. Individual F_2_ progeny were placed onto separate NGM dishes and F_3_ progeny were genotyped to identify animals homozygous for *raga-1(viz128)* and *depd-5(am346)*. The *depd-5(am346)* genotyping assay is described above.

*raga-1* Q63L FPe: TCTGATGGGCAAAAGTGGAT
*raga-1* Q63L FPi: GATTGTGGTGGTCTTGAAAGT
*raga-1* Q63L RPe: GACGTTTGGCACCAGATGAT
*raga-1* Q63L RPi: AAGTTCTCCATGAATGATTCCT

The FPe and RPe amplify a 608 bp amplicon in every genotype while the FPe and RPi make a 241 bp wild-type amplicon and the FPi and RPe generate the 402 bp *viz128* amplicon. The cycling conditions were modified to include a 63°C annealing temperature and a 45 seconds elongation time.

### Construction of the raga-1(ok386) II; depd-5(am346) III double mutant

The *raga-1(ok386) II; depd-5(am346) III* double mutant was constructed as described for double mutant above. The *depd-5(am346)* genotyping assay is described above. The F_3_ progeny from the genetic cross were genotyped using these primers to identify animals homozygous for *raga-1(ok386)*.

*raga-1(ok386)* F1: GATCCGGCAAGACTTCGATG
*raga-1(ok386)* F2: CCGCATTCATCGACTCGTTC
*raga-1(ok386)* R: AAGGATCGGAACTGAAGCGA

The *raga-1(ok386)* F1 and R primers produce a 1,535 bp amplicon on the wild-type *raga-1* locus and a 243 bp amplicon in the *raga-1(ok386)* mutant. The *raga-1(ok386)* F2 and R primers produce a 671 bp amplicon on the wild-type *raga-1* locus and no product in the *raga-1(ok386)* mutant.

### Construction of the szt-2(gk554651) II; depd-5(am346) III double mutant

The *szt-2(gk554651) II; depd-5(am346) III* double mutant was constructed as described for the double mutants above. The *depd-5(am346)* genotyping assay is described above. The F_3_ progeny from the genetic cross were genotyped using these primers to identify animals homozygous for *szt-2(gk554651)*.

*szt-2(gk554651)* F: CACATCGTCACTCCAGCAAC
*szt-2(gk554651)* R: AACTCACCGCCATAACAACG

The *szt-2(gk554651)* F and R primers produce a 1,268 bp amplicon for both the wild-type and gk554651 mutant alleles. The PCR product was digested with BsrI yielding two restriction fragments, 804 and 464 bp, from the gk554651 allele.

### Construction of the tph-1(ot1545) II; depd-5(am346) III double mutant

The *tph-1(ot1545) II; depd-5(am346) III* double mutant was constructed as described for the double mutants above. The *depd-5(am346)* genotyping assay is described above. The F_3_ progeny from the genetic cross were genotyped using these primers to identify animals homozygous for *tph-1(ot1545)*.

tph-1(ot1545) F1: TCCCACGGAAACCATGACAG
tph-1(ot1545) F2: GGTCACATGGCTCTATTCGC
tph-1(ot1545) R: ACCTGCCCGACATGTTAGAG

The *tph-1(ot1545)* F1 and R primers produce a 3,262 bp amplicon from the wild-type *tph-1* locus and a 390 bp amplicon in the *tph-1(ot1545)* mutant. The *tph-1(ot1545)* F2 and R primers produce a 1,229 bp amplicon on the wild-type *tph-1* locus and no product in the *tph-1(ot1545)* mutant.

### Construction of the depd-5(am346) III tdo-2(zg218) III double mutant

The *depd-5(am346) III tdo-2(zg218) III* double mutant was constructed as described for the double mutants above. The F_3_ progeny from the genetic cross were genotyped using these primers to identify animals carrying the wild-type *tdo-2* allele and to screen for candidate *tdo-2(zg218*) homozygotes based on the absence of amplification.

*tdo-2(zg218)* F: GACACAATGAGTCCATTGGA
*tdo-2(zg218)* R: ACCGGTTCCTTGTTTAGCAC
tdo-2(zg218) F(seq): CCGATGGTCAACGTGTAGAC

The *tdo-2(zg218)* F and R primers produce a 1,403 bp amplicon from the wild-type *tdo-2* locus and no amplicon in the *tdo-2(zg218)* mutant. The *tdo-2(zg218)* F(seq) and R primers were used to generate an amplicon utilized for Sanger sequencing (described later).

Because the *depd-5* and *tdo-2* loci are genetically linked, identification of the desired recombinant required high-throughput screening. Pyrosequencing was performed using model and the PyroMark Q48 Advanced Reagents (4×48) kit (Qiagen, #974002).

*depd-5(am346)* F: ATGGAAGATGATCGTAAACACTC
*depd-5(am346)* Sequencing F: AAAAGTACAAAGTATGGG
*depd-5(am346)* Biotinylated R: GGGAAAGAAATCAGTTGTAATTGG
*depd-5(am346)* G(wt)>A
ATGGAAGATGATCGTAAACACTCTTCAAAATCGAATTCCGCTGGAAATGCGAGTAAAAAGTACAAAGTATGGGCATG[G/A]GGATCAACTGGAGAAGAGAAATGGAATGTTGATATGGAAATTGGAACCGATTGGAAATCACTTGTCAGATCTGCATTACTTCCAATTACAACTGATTTCTTTCCC

Sanger confirmation was especially important for *tdo-2(zg218)* because failure to amplify could reflect either the deletion allele or technical failure of the PCR. Therefore, candidate recombinants identified by PCR and pyrosequencing were confirmed by Sanger sequencing of both loci.

### Construction of the *depd-5(am346) III; aak-2(ok524) X* double mutant

The *depd-5(am346) III; aak-2(ok524) X* double mutant was constructed as described for the double mutants above. The *depd-5(am346)* genotyping assay is described above. The F_3_ progeny from the genetic cross were genotyped using these primers to identify animals homozygous for *aak-2(ok524)*.

*aak-2(ok524)* F1: AAGGAGACACTCGGAGTTGG

*aak-2(ok524)* F2: TCATTTCCGGCGTTGACTAC

*aak-2(ok524)* R: TTCATCGGGTCTACGCAGAG

The *aak-2(ok524)* F1 and R primers produce a 1,284 bp amplicon from the wild-type *aak-2* locus and a 876 bp amplicon in the *aak-2(ok524)* mutant. The *aak-2(ok524)* F2 and R primers produce a 678 bp amplicon on the wild-type *aak-2* locus and no product in the *aak-2(ok524)* mutant.

### Construction of the *depd-5(am346) arf-1(ok796)* III double mutant

The *depd-5(am346) arf-1(ok796)* double mutant was constructed as described for the double mutants above. The F_3_ progeny from the genetic cross were genotyped using these primers to identify animals carrying the wild-type *arf-1* allele and to screen for candidate *arf-1(ok796)* homozygotes based on the absence of amplification.

*arf-1(ok796)* F1: ATGGTTGGATTGGATGCTGC
*arf-1(ok796)* F2: TGATGCCGTTCTTCTGGTCT
*arf-1(ok796)* R: ACGTGAACACTGAGAATGCG

The *arf-1(ok796)* F1 and R primers produce an 1,836 bp amplicon from the wild-type *arf-1* locus and a 696 bp amplicon in the *arf-1(ok796)* mutant. The *arf-1(ok796)* F2 and R primers produce a 1,377 bp amplicon on the wild-type arf-1 locus and no product in the *arf-1(ok796)* mutant.

Because the *depd-5* and *arf-1* loci are genetically linked, identification of the desired recombinant required high-throughput screening. Pyrosequencing was performed as described above. Candidate recombinants identified by PCR and pyrosequencing were confirmed by Sanger sequencing of both loci.

### Generation of Transgenic Strains

#### Construction of WU1940: the *hlh-30* Deletion in CB4856

The Left and Right CRISPR sgRNA plasmids were generated by ligating the paired sgRNA oligonucleotides into the U6 [3055] vector using a 1 x SAP Reaction Buffer, 50 ng of U6 [3055], sgRNA F and R (IDT) in an equimolar ratio of 0.5 ng/μl, 10 units of Bsa-1 (New England Biolabs), 100 units of T4 DNA Ligase (New England Biolabs) in 37°C 15 min, three cycles of 15°C 5 min and 37°C 10 min, then 55°C 5 min and 80°C 5 min. The plasmids were transformed into Top 10 Chemically Competent cells (Invitrogen) and bacterial colonies were selected on LB Kanamycin dishes.

Left sgRNA F: TTTG CG CAA AAT CGA CCG AAA GT
Left sgRNA R: AAAC AC TTT CCG TCG ATT TTG CG
Right sgRNA F: TTTG GG ACA CTC TGA CAC CAC GT
Right sgRNA R: AAAC AC GTG GTG TCA GAG TGT CC
*dpy-10(cn64)* self-repair template: ACT TGA ACT TCA ATA CGG CAA GAT GAG AAT GAC TGG AAA CCG TAC CGC ATG CGG TGC CTA TGG TAG CGG AGC TTC ACA TGG CTT CAG ACC AAC AGC CTA T
*hlh-30* deletion self-repair template: ACC GGT TTC TCA TCA GAT CCT CCT CCT ACT CGT AGG TTT TGT ATA GTT TTT GGC GGG AAG

Injected CRISPR reagents into CB4856 young adults, then placed them at 25°. Reagents included: 600 nM *dpy-10(cn64)*, 20 ng/μl *dpy-10* sgRNA [3153], 600 nM *hlh-30* deletion self-repair template, 50 ng/μl left sgRNA [3055], 50 ng/μl right sgRNA [3055] and 50 ng/μl Cas9 [3143]. Selected F_1_ rollers, F_2_ rollers and F_3_ non-roller (wild type) worms. Detected 692 bp deletion spanning exons 5 and 6 with PCR and confirmed with Sanger sequencing. The resulting allele was designated *am396*.

#### Construction of WU2001 to restore the *am346* Lesion

The reversion CRISPR sgRNA plasmid was generated by ligating the forward and reverse reversion sgRNA oligonucleotides U6 [3055] vector.

Reversion sgRNA F: TTT GTA TGG GCA TGA GGA TCA AC
Reversion sgRNA R: AAA CGT TGA TCC TCA TGC CCA TA
Reversion Repair Template: CGC TGG AAA TGC GAG TAA AAA GTA CAA AGT ATG GGC GTG GGG ATC CAC TGG AGA AGA GAA ATG GAA TGT TGA TAT GGA AAT T

Young adult *depd-5(am346)* hermaphrodites were injected with CRISPR reagents, then placed at 20°C. The injection mixture contained: 600 nM *dpy-10(cn64)* repair template, 20 ng/μL *dpy-10* sgRNA plasmid [3153], 600 nM reversion repair template, 50 ng/μL reversion sgRNA and 50 ng/μL Cas9 [3143]. Selected F_1_ roller progeny and placed one per NGM dish. After 24 hours of egg laying, prepared DNA from the single adult F1 worm using ThermoPol Buffer (B9004S NEB Ipswich, MA), Proteinase K (69504 Qiagen) and water. Digested 60°C for 60 min and 95°C for 15 min. Pulsed and centrifuged. PCR with Taq Plus Master Mix Red 2 x D124R (Lamda Biotech, St. Louis MO) with 0.4uM T08A11.1 F and R primers, 1uL worm DNA in a 25uL reaction. 95°C 5 minutes, 30 cycles of 94°C 30 seconds, 64°C 30 seconds and 72°C for 90 seconds, and 72°C for 5 minutes. *depd-5* F: ATA CGC ACT CAC TTC GGA TG and *depd-5 R*: TGA GCC ATC GTA GGC TTC TC produce a 1,256 bp amplimer. Presence of CRISPR-modified DNA introduces BamHI restriction site and BamHI digestion produces 857 and 399 base pair restriction fragments. Identified worm that was heterozygous for the CRISPR modification. Placed individual F2 worms from its NGM dish, onto 30 new NGM dishes. Identified dishes containing worms that were homozygous for the CRISPR modification with DNA from approximately 10 F3 worms, and the same approach of PCR and analysis of BamHI restriction fragments. Verified the homozygous CRISPR modification with Sanger sequencing (Azenta Life Sciences, Chelmsford, MA).

### EMS Mutagenesis

To isolate mutants that could not be rescued with glucose and linoleic acid following starvation, we mutagenized *hlh-30(tm1978)* hermaphrodites with ethyl methanesulfonate (EMS) as described.^39^ We picked mutagenized *hlh-30* hermaphrodites at the L4 larval stage and allowed these animals to self-fertilize on NGM dishes. Visibly healthy F_1_ progeny were placed, one animal per dish, on NGM dishes and allowed to lay progeny. One visibly healthy F_2_ animal from each F_1_ dish was placed on an NGM dish and allowed to self. Immediately before bleaching the population of gravid F_3_ adults, one F_3_ was moved to a fresh NGM dish to ensure survival of the strain. The remaining adult hermaphrodite F_3_ progeny were treated with alkaline hypochlorite solution and F_4_ eggs were washed and resuspended in M9 at a density equal or less than four to six worms per 1μL. The L1s of the F_4_ generation were starved for 33 hours, then transferred to CeMM containing glucose and linoleic acid for 15 hours, then placed on NGM dishes seeded with OP50. Dishes were screened for worms that failed to recover and given an allele designation.

### Whole Genome Sequencing and Mapping of Mutants

The *am346* mutant was outcrossed to the *hlh-30(tm1978)* strain twice through testing under the same conditions of the screen and selection of F_4_ progeny whose siblings failed to recover. The outcrossed Bristol *am346* hermaphrodite was crossed to the Hawaiian *hlh-30(am396)*^40^ male, the progeny were tested using the same conditions of the screen, and progeny of F_2_ recombinants that failed to recover were kept and retested to yield 36 recombinants whose DNA was pooled and submitted for whole genome library prep, NovaSeq S4 2×150 (4 lanes) whole-genome sequencing and Hawaiian SNP mapping as described.^13^

### Mapping of Mutants

Whole genome sequencing was performed by the Washington University in St. louis Genome Technology Access Center (GTAC). The WS220.64 C. elegans reference genome, as well as a list of Hawaiian SNPs, were obtained from sourceforge.net/projects/mimodd/files/MiModD-0.1.9/. Analyses were performed using the MiModD v0.1.9 toolkit and the usegalaxy.org platform. Whole genome sequencing reads were aligned to the WS220.64 reference genome using the bowtie2 read-alignment tool.^41^ Next, the MiModD Variant Calling tool was used to generate a list of variants that differed from the reference genome; these variants result either from EMS mutagenesis or from the Hawaiian background. We then used the MiModD Extract Variant Sites and MiModD NacreousMap tool to locate regions of the genome that contained no Hawaiian SNPs; these regions are tightly linked to the original *am346* strain, and therefore are likely to contain the causative mutation

We identified a region on Chromosome III from 4.5-10 Mb that contained no Hawaiian SNPs. We used the MiModD VCF Filter tool to generate a list of the 212 variants found in this region. We excluded small insertion and deletion (indel) variants; these are likely sequencing errors, as EMS mutagenesis tends to produce base pair changes. We next used the UCSC Genome Browser to analyze each variant and its surrounding regions. RefSeq display was used to determine if each variant was in a known *C. elegans* gene, an intergenic region, or in a 3’ untranslated region (UTR). If a variant was in a gene-encoding region, the transcript for the gene was copied from Wormbase. Then, using A plasmid Editor (ApE), the sequence containing the variant (10 nucleotides upstream and downstream of the SNP) was searched for in the transcript and the reading frame was determined. Once the variant’s position in the reading frame was found, the nature of the mutation was determined using an amino acid/codon chart.

We identified a total of 212 variants (**Table S2**). 94 were in intergenic regions, 84 were in introns, 9 were silent mutations, 3 were in 3’ UTRs, 2 were in a non-coding transcript (K04H4.13), and 9 were due to disagreements between the UCSC and Wormbase reference sequences (this was determined by searching for the mutated/variant sequence in the transcript, as opposed to the reference sequence). The remaining 11 variants were non-silent mutations in gene-encoding regions (**Table S3**).

### Preparation of CeMM Formulations

The composition of all CeMM formulations used in this study is provided in **Table S1**. Glassware was soaked overnight in a 1% solution of detergent (company), then rinsed three times with distilled water, soaked in 0.18 M HCl overnight, rinsed with distilled water three times, dried and autoclaved. CeMM was prepared as described.^9,12^

Nutrient omission and supplementation experiments were generated by preparing CeMM lacking specific nutrient classes or by adding individual nutrients at the concentrations indicated in Table 3 (only made Minimal, or with nucleic acids, amino acids, essential, or nonessential, or individual or -tryptophan, never made with glucose or with linoleic acid because this was added on the day of the experiment. Then filter sterilized 0.22 μm filter, wrapped in aluminum foil to protect against light and stored at 4 degrees. Water used in CeMM formulations was purified with the Milli-Q Synthesis A10 system purification system equipped with a BioPak polisher (CDUFBI001, MilliporeSigma, Burlington, MA, USA).

Vehicle control preparations were generated by replacing linoleic acid with an equivalent volume of ethanol and water while mainlining identical concentrations of NaOH and BSA. For example, the BSA conjugated control solution contained 7.85mL filtered water, 150 μL 1 M NaOH, 45 μL 100% ethanol and 45 μL additional water, which was then added to 9.9 mL of 20% BSA.

### Starvation and Recovery Assays

For all assays reported, 50-120 animals were studied per biological replicate, except as indicated. For bleaching, gravid adults fed for at least the past three generations were transferred from *OP50* seeded NGM dishes to 15 mL tubes (MidSci name) using water (machine name) and glass Pasteur pipets. The tubes were spun in centrifuge (name) at 400 x g for one minute, then the supernatant was aspirated leaving 0.1 mL of liquid and the worm pellet. The worm pellet was washed by resuspending in water up to the 14.5 mL mark on the tube, inverted 10 times to mix, then spun at 400 x g for one minute. The supernatant was aspirated leaving 0.1 mL of liquid completing the wash. The worm pellet was washed two additional times. Alkaline hypochlorite was made immediately before addition to the worms using these a 2:3 ratio of 4 M NaOH and freshly-made 20% bleach. Alkaline hypochlorite was inverted 10 times to mix, then 5 mL was added to the tubes containing worm pellets. Worm suspensions were exposed to alkaline hypochlorite for 4–5 min, with continuous inversion, until adult carcasses were dissolved and embryos remained visible. Following bleaching, samples were diluted to 14.5 mL with water and centrifuged immediately. The tubes were inverted to mix five times, then immediately spun at 1,000 x g for two minutes. The supernatant was aspirated leaving 0.1 mL of liquid and the worm pellet. Three washes were conducted as before using the new centrifugation speed. After the third wash, the worm pellet was resuspended with M9 and the density of eggs was adjusted to four to six eggs per uL. The final volume of M9 inside the tube varied across experiments but was always between eight and 12 mL. Tubes were laid horizontally on a shaker (company) rotating 100 rpm inside a 20°C incubator.

Preparation of CeMM minimal solution is described in **Table S1**. For preparing supplements to CeMM minimal, additives such as glucose and linoleic acid was added to CeMM minimal upon the day of the experiment. Glucose was added at 180 mM final concentration in 1 x CeMM. CeMM containing linoleic acid (Cayman Chemical 90150-500mg) was made using four steps. First, the contents of the linoleic acid bottle which was approximately 500 μL were transferred to a 1.7 mL Eppendorf tube along with the addition of 500 μL of 100% ethanol to bring the volume up to 1 mL to create a 1.78 M solution. Second, a 20% solution of fatty-acid free bovine serum albumin solution (Lampire Biological Laboratories #7500804) was prepared by gently layering BSA on the surface of 1 x PBS where it dissolved overnight. The next day it was filter sterilized with a 0.22 μM filter. Third, a 20 mM solution of linoleic acid was prepared by adding 90 μL of a 1.78 M linoleic acid stock solution to a tube containing 7.85 mL water previously placed in 72°C water bath for 30 minutes. This cloudy solution of 20 mM linoleic acid was made soluble through the addition of 150 μL of 1 M NaOH, in 50 μL aliquots with vortexing after each addition. Fourth, conjugation of linoleic acid to BSA was accomplished when 6 mL of 20 mM Linoleic acid was added to 9.9 mL 20% BSA, immediately pipetted to mix creating a 7.55 mM solution of linoleic acid. Glucose added to solutions without filtering.

Preparation of CeMM final compounding of solutions to make 1 x CeMM with a final concentration of 1 mM linoleic acid was as follows: added 1.375 mL conjugated linoleic acid BSA, 3.625 mL water and 5 mL 2X CeMM. Placed in 1.7 mL eppendorf tubes at a final volume of 1 mL 1X CeMM. The tube holding worms starved for 24 hours or the designated time, would be spun at 1,000 x g and the supernatant aspirated to leave 0.5 mL. Then 50 μL would be transferred 1 mL of 1XCeMM. The worms would be returned to the horizontal shaker in the 20°C incubator for 24 hours.

Plating the worms on NGM dishes involved centrifugation of the tube, followed by aspirating all but 100 uL, then pipetting this liquid onto the OP50 found on the NGM dish. Alive worms were counted as moving worms. Dishes were returned to the 20°C incubator. Two days later the worms were scored as recovered if they were alive, then the number of recovered worms divided by the initial number of alive worms on that dish was reported as Recovery (%).

### Targeted Metabolomics

Animals were prepared using the starvation and recovery assay described above. Samples were prepared following either 24 hours or seven days of starvation in M9. For the group starved seven days, the M9 medium was replaced through centrifugation of animals, aspiration of theold M9 and resuspension in fresh M9, after two and six days of starvation. Following starvation, animals were incubated for 24 hours in CeMM minimal CeMM, or CeMM minimal containing glucose or CeMM minimal containing glucose and tryptophan. Following the 24-hour CeMM incubation, animals were collected and washed twice with M9. After the final wash, as much M9 as possible was removed and the worm pellets were snap frozen in 1.7 mL Eppendorf tubes in liquid nitrogen and stored at -80°C until analysis.

To prepare samples, worm pellets were resuspended in 80% (v/v) methanol and tip-sonicated for 3 cycles (30 s ON, 59 s OFF), followed by vigorous vortexing for 1 min and overnight incubation at -80 °C. Heavy isotope-labeled internal standards (Trp-d5, Kyn-d4, and 5HT-d4) were added to each sample at a final concentration of 12.5 ng/mL. Samples were centrifuged at 21,000 × g for 15 min at 4 °C, and the supernatants were collected and dried completely in a SpeedVac without heat. For derivatization, dried extracts were reconstituted in 25 μL of 100 mM sodium bicarbonate buffer (pH 10.5), mixed with 25 μL of 1 mg/mL dansyl chloride, and incubated at 60 °C for 5 min. After cooling to room temperature, samples were centrifuged at 21,000 × g for 10 min at 4 °C, and the supernatants were transferred to amber glass vials for LC-MS/MS analysis.

For Liquid Chromatography-Mass Spectrometry (LC-MS/MS) analysis, chromatographic separation was performed using a Waters ACQUITY UPLC HSS T3 C18 column (2.1 × 150 mm, 1.8 μm) with water containing 0.1% formic acid as mobile phase A and 90% acetonitrile containing 0.1% formic acid as mobile phase B. The flow rate was 0.4 mL/min, the injection volume was 5 μL, and the column temperature was maintained at 50 °C. The 10-min gradient was: 0-0.7 min, 0% B; 2 min, 0.5% B; 6-8 min, 98% B; and 8.1-10 min, 0% B. MS analysis was performed on an Orbitrap Exploris 240 mass spectrometer equipped with a HESI-II source operated in positive ion mode. Target compounds were analyzed by parallel reaction monitoring (PRM) at 15,000 resolution using a 2.0 m/z isolation window, RF lens setting of 70%, and automatic AGC target and maximum injection time. Collision energies were optimized individually for each compound. Quantifier product ions were m/z 159.0913 for dansyl-Trp, 120.0442 for dansyl-Kyn, and 146.0599 for dansyl-5HT; the corresponding heavy internal-standard quantifier ions were m/z 164.1227, 122.0567, and 148.0724, respectively.

Absolute quantification was performed as follows. Calibration curves for tryptophan, kynurenine, and serotonin were prepared by serial dilution of dansyl chloride-derivatized analytical standards in wild-type worm extract and analyzed in triplicate. The calibration range was 95 pg/mL to 12.5 μg/mL for each analyte. Linear ranges were defined by R² > 0.99 and coefficient of variation (CV) < 20%, and the lower limit of quantification (LLOQ) was defined as the lowest concentration within the linear range above the limit of detection (LOD), with the LOD determined as three standard deviations of the blank signal. Absolute concentrations were determined from analyte-to-heavy internal standard peak area ratios using linear regression of the corresponding calibration curves. Metabolite abundance was normalized to the number of animals in each sample and reported as pg per 1,000 worms.

### Generation of DEPDC5 mutant HeLa cells

The *DEPDC5* gene was targeted with CRISPR-Cas9 based strategy in a HeLa cell line with TFEB-GFP expression,^42^ at a location analogous to that in *depd5(am346)* worms by the Genome Engineering & Stem Cell Center (GESC@MGI) at Washington University in St. Louis. Briefly, gRNAs were designed to cleave the *DEPDC5* gene so that an out-of-frame indel leads to premature stop codons in between the SABA and SHEN domains. Synthetic gRNA with target sequence 5’-GGGCAGTGTGCGGCTTGGTA was ordered from Integrated DNA Technologies (Coralville, IA) and complexed with Cas9 protein before delivered into the parental cells using Nucleofection. The transfected cells were then single cell sorted into 96-well plates, and single cell clones were screened using NGS for harboring only out-of-frame alleles. WT and DEPDC5-mutant TFEB-GFP HeLa cells were maintained under fed conditions or subjected to starvation in Dulbecco’s phosphate-buffered saline (DPBS) containing Mg²⁺ and Ca²⁺ (Gibco, cat#14040-133) for 0.5, 1.0 or 1.5 hours. Starvation treatments were initiated sequentially, beginning with the 1.5-hour group, followed by the 1-hour and 0.5-hour groups, to ensure that all samples were collected simultaneously at a common endpoint. Fed control cells were maintained in DMEM (Cat# 10013CV, Corning™), supplemented with 10% inactivated FBS (Cat# 16140071, Gibco), 2 mM L-glutamine (Cat# 25030081, Life Technologies), and Penicillin/streptomycin (100 U/mL) (Cat# 15140122, Life Technologies).

### Immunoblotting

Immunoblotting was performed on cellular extracts as previously described.^43^ Specific primary antibodies employed are as follows: p-mTOR (Ser2448) (Cell Signaling, 5536S), mTOR (Cell Signaling, 2972S), p-p70 S6K (Thr389) (Cell Signaling, 9234S), p70 S6K (Cell Signaling, 2708S), p-4E-BP1 (Thr37/46) (Cell Signaling, 2855S), 4E-BP1 (Cell Signaling, 9644S), DEPDC5 (Abcam, ab213181). The following secondary antibodies were used for immunoblotting: Anti-rabbit IgG HRP-linked (Cell Signaling, 7074S).

### Analysis of TFEB nuclear localization

TFEB nuclear localization was quantified at the single-cell level using CellProfiler software with an image-analysis pipeline as described.^42^ Nuclei were segmented from DAPI images, and TFEB-GFP-positive cells were identified from the GFP channel. Nuclear TFEB-GFP fluorescence was measured within the segmented nuclear region, whereas cytoplasmic fluorescence was quantified within a 10-pixel-wide perinuclear region extending outward from the nuclear boundary. For each cell, a TFEB nuclear translocation (NT) index was calculated as N/(N+C), where (N) and (C) represent the GFP fluorescence intensities measured in the nuclear and perinuclear cytoplasmic compartments, respectively. The NT index ranges from 0 to 1, with higher values indicating greater nuclear localization of TFEB-GFP. Single-cell NT index values were displayed as violin plots, with approximately 100-180 cells analyzed per group.

### Statistics

All data are presented as mean ± SEM (standard error of the mean). All measurements were obtained on distinct biological replicates. Statistics were performed in Prism Version 9.1.2 (GraphPad Software Inc.). Data were tested for assumptions of normality with Shapiro-Wilk normality test. Statistical significance of differences was calculated via unpaired 2-tailed Student’s t test for 2 group comparisons, or one-way analysis of variance (ANOVA) for assessing differences across multiple groups followed by post-hoc testing to evaluate pairwise differences. A non-parametric test was employed if data were not normally distributed as indicated in the figure legend. In specific instances where analysis of variance was performed, data were subjected to transformation as indicated and re-assessed for normality if they failed normality testing, prior to statistical testing. A *P* value < 0.05 considered statistically significant. The datasets from the current study are available from the corresponding author on reasonable request.

## Author contributions

J.T.M., R.C., D.S., S.K., H.Z., A.D., M.G., M.I., K.R.Q.L., D.L.S., X.M., and B.M.E. performed experiments, acquired, and analyzed the data; J.T.M and D.R.R. interpreted the data; and J.T.M, K.K. and A.D. conceived the experiments, analyzed the data, supervised the work, and drafted the manuscript.

## Acknowledgements

We thank the Mass Spectrometry Technology Access Center (MTAC@MGI) at Washington University in St. Louis for metabolite assays. The Mass Spectrometry Technology Access Center at the McDonnell Genome Institute (MTAC@MGI) at Washington University School of Medicine is supported by the Diabetes Research Center/NIH grant P30 DK020579, Institute of Clinical and Translational Sciences/NCATS CTSA award UL1 TR002345, and Siteman Cancer Center/NCI CCSG grant P30 CA091842. Some strains were provided by the CGC, which is funded by NIH Office of Research Infrastructure Programs (P40 OD010440). We thank Marco Sardiello, Ph.D. for providing the TFEB-GFP expressing HeLa cells. We also thank the Genome Engineering & Stem Cell Center at Washington University in St. Louis for creating the KO cell lines.

