## Supplementary Data for "Tryptophan Restores Recovery Competence Following Starvation in the Absence of Lysosomal Amino Acid Sensing"

**Supplementary Table S1: Table of CeMM constituents**

| **CeMM Minimal** | | | |
|---|---|---|---|
| Component | Final Concentration (2X) | Supplier | Catalog Number |
| **Dry Ingredients** | | | |
| Choline H2 citrate | 0.6 mM | Millipore-Sigma | C2004 |
| Glutathione | 1.33 mM | Millipore-Sigma | G6529 |
| Myoinositol | 0.722 mM | Millipore-Sigma | I5125 |
| Cytochrome C | 0.00807 mM | Millipore-Sigma | C7752 |
| **Salt Solution** | | | |
| citric acid monohydrate | 6 mM | Millipore-Sigma | C0706 |
| magnesium hydroxide | 6 mM | Millipore-Sigma | M5421 |
| potassium citrate tribasic monohydrate | 3 mM | Millipore-Sigma | P1722 |
| potassium phosphate monobasic | 18 mM | Millipore-Sigma | P5655 |
| ferrous ammonium sulfate hexahydrate | 0.3 mM | Millipore-Sigma | F1543 |
| calcium chloride dihydrate | 3 mM | Millipore-Sigma | C5080 |
| copper chloride dihydrate | 0.075 mM | Millipore-Sigma | 307483 |
| manganese chloride tetrahydrate | 0.225 mM | Millipore-Sigma | M8054 |
| zinc chloride | 0.150 mM | Millipore-Sigma | Z0152 |
| **TEA-Soluble Vitamins & Growth Factors** | | | |
| p-aminobenzoic acid | 109.5 µM | Millipore-Sigma | A9878 |
| Biotin | 30.7 µM | Millipore-Sigma | B4639 |
| Folic acid | 34 µM | Thermo Scientific Chemicals | 216630100 |
| Lipoic acid | 36.4 µM | Millipore-Sigma | |
| Niacin | 121.8 µM | Millipore-Sigma | N0761 |
| **Water-Soluble Vitamins & Growth Factors** | | | |
| n-acetyl-D-glucosamine | 135.6 µM | Millipore-Sigma | A8625 |
| Cyanocobalamin | 5.5 µM | Millipore-Sigma | V6629 |
| Niacinamide | 122.8 µM | Millipore-Sigma | N3376 |
| D-pantethine | 13.5 µM | Millipore-Sigma | P2125 |
| D-pantothenic acid | 62.9 µM | Millipore-Sigma | P5710 |
| Pyridoxal monophosphate | 30.4 µM | Millipore-Sigma | P3657 |
| Pyridoxamine dihydrochloride | 31.1 µM | Millipore-Sigma | P9158 |
| Pyridoxine hydrochloride | 73 µM | Millipore-Sigma | P6280 |
| Riboflavin 5'-phosphate | 29.2 µM | Millipore-Sigma | R7774 |

| | | | |
|---|---|---|---|
| Thiamine hydrochloride | 44.5 µM | Millipore-Sigma | T1270 |
| **Additives to CeMM Minimal** | | | |
| **Essential Amino Acids** | | | |
| L-arginine | 10 mM | Millipore-Sigma | A8094 |
| L-histidine | 4 mM | Millipore-Sigma | H8000 |
| L-isoleucine | 12 mM | Millipore-Sigma | I7403 |
| L-leucine | 20 mM | Millipore-Sigma | L8912 |
| L-lysine-HCl | 15 mM | Millipore-Sigma | L5626 |
| L-methionine | 5 mM | Millipore-Sigma | M5308 |
| L-phenylalanine | 8 mM | Millipore-Sigma | P5482 |
| L-threonine | 12 mM | Millipore-Sigma | T8441 |
| L-tryptophan | 2 mM | Millipore-Sigma | T8941 |
| L-valine | 20 mM | Millipore-Sigma | V0513 |
| **Nonessential Amino Acids** | | | |
| L-alanine | 30 mM | Millipore-Sigma | A7469 |
| L-aspartic acid | 25 mM | Millipore-Sigma | A8949 |
| L-cysteine-HCl-$H_20$ | 3 mM | Millipore-Sigma | C6852 |
| L-glutamate(Na) $H_20$ | 6 mM | Millipore-Sigma | G1626 |
| L-glutamine | 20 mM | Millipore-Sigma | G3126 |
| glycine | 20 mM | Millipore-Sigma | G7403 |
| L-proline | 10 mM | Millipore-Sigma | P0380 |
| L-serine | 15 mM | Millipore-Sigma | S4500 |
| L-tyrosine | 3 mM | Millipore-Sigma | T8566 |
| **Carbohydrates** | | | |
| D-Glucose | 361 mM | Millipore-Sigma | G8270 |
| **Lipids** | | | |
| Linoleic Acid | 2mM | Cayman Chemical | 90150 |
| **Nucleic Acids** | | | |
| adenosine 5'-monophosphate sodium salt | 2 mM | Millipore-Sigma | A1752 |
| cytidine 5'-monophosphate disodium salt | 2 mM | Millipore-Sigma | C1006 |
| guanosine 5'-monophosphate disodium salt hydrate | 2 mM | Millipore-Sigma | G8377 |
| thymine | 2 mM | Millipore-Sigma | T0895 |
| uridine 5'-monophosphate disodium salt | 2 mM | Millipore-Sigma | U6375 |

**Table S2. Variant Filtering and Candidate Coding Mutations Identified following CB4856 Mapping and Whole Genome Sequencing**

| Category | Number of Variants |
|---|---|
| **Total Variants Identified** | **212** |
| Remove Intergenic Variants | 94 |
| Remove Intronic Variants | 84 |
| Remove Silent Mutations | 9 |
| Remove 3' UTR Variants | 3 |
| Remove Non-Coding Transcript Variants | 2 |
| Remove Reference Discrepancies | 9 |
| **Variants causing nonsense or missense mutations** | **11** |

Reference sequence discrepancies were identified by comparison of UCSC and WormBase reference sequences. Candidate variants were evaluated by searching the mutant sequence withing the transcript using ApE.

**Table S3. Coding Candidate Mutations Identified Following CB4856 Mapping and Whole Genome Sequencing**

| Gene | Putative Mutation |
|---|---|
| F25F2.1 | L215F |
| dpf-4 | S45L |
| ubq-1 | I183N, I213T or I441T |
| nono-1 | P502Q, K503N, S504P |
| C09E7.10 | R94STOP |
| B0303.7 | C16S |
| cls-3 | V159I |
| mcm-6 | S568N |
| T08A11.1 | W917STOP |

**Table S4: Table of *C. elegans* strains used in this study**

| Strain | Genotype | Common Name | Source | Notes |
|---|---|---|---|---|
| **WT and Parental Strains** | | | | |
| N2 | Wildtype | WT | CGC | |
| JIN1375 | *hlh-30(tm1978)* | *hlh-30(lf)* | CGC | EMS parent strain |
| WU1997 | *depd-5(am346)* | *depd-5(lf)* | this study | |
| WU1998 | *depd-5(gk657858) III* | | CGC | Q536STOP |
| WU2185 | *depd-5(ve878) III* | | CGC | Δ2-7 |
| AUM1693 | *raga-1(viz128) II* | *raga-1(gf)* | Arur lab | Q63L |
| WU2172 | *raga-1(ok386) II* | *raga-1(lf)* | CGC | |
| WU2183 | *szt-1(gk554651) II* | *szt-2(lf)* | CGC | |
| OH19337 | *tph-1(ot1545) II* | *tph-1(lf)* | CGC | |
| OW717 | *tdo-2(zg218) III* | *tdo-2(lf)* | CGC | |
| RB754 | *aak-2(ok524) X* | *aak-2(lf)* | CGC | |
| VC567 | *arf-1(ok796) III* | *arf-1(lf)* | CGC | |
| **EMS-Derived Strains** | | | | |
| WU1880 | *hlh-30(tm1978) II; depd-5(am346) III* | *hlh-30(lf); depd-5(lf)* | this study | W917STOP |
| **Constructed Double Mutants** | | | | |
| WU2181 | *raga-1(viz128) II; depd-5(am346) III* | *raga-1(gf); depd-5(lf)* | this study | |
| WU2182 | *raga-1(ok386) II; depd-5(am346) III* | *raga-1(lf); depd-5(lf)* | this study | |
| WU2184 | *szt-2(lf) II; depd-5(am346) III* | *szt-2(lf); depd-5(lf)* | this study | |
| WU2247 | *tph-1(ot1545) II; depd-5(am346) III* | *tph-1(lf); depd-5(lf)* | this study | |
| WU2251 | *tdo-2(zg218) III depd-5(am346) III* | *tdo-2(lf) depd-5(lf)* | this study | |
| WU2303 | *depd-5(am346) III; aak-2(ok524)* | *depd-5(lf); aak-2(lf)* | this study | |
| **Transgenic Strains** | | | | |
| WU1940 | *hlh-30(am396) II* | | this study | Hawaiian strain |
| WU1999 | *depd-5(am397) III* | *HA-DEPD-5* | This study | |
| WU2000 | *depd-5(am398) III* | *HA-depd-5(lf)* | This study | |

| | | | | |
|---|---|---|---|---|
| WU2001 | *depd-5(am399) III* | *reversion mutant* | This study | STOP917W |

Indicated strains were provided by the CGC, which is funded by NIH Office of Research Infrastructure Programs (P40 OD010440).

**Supplementary Figures**

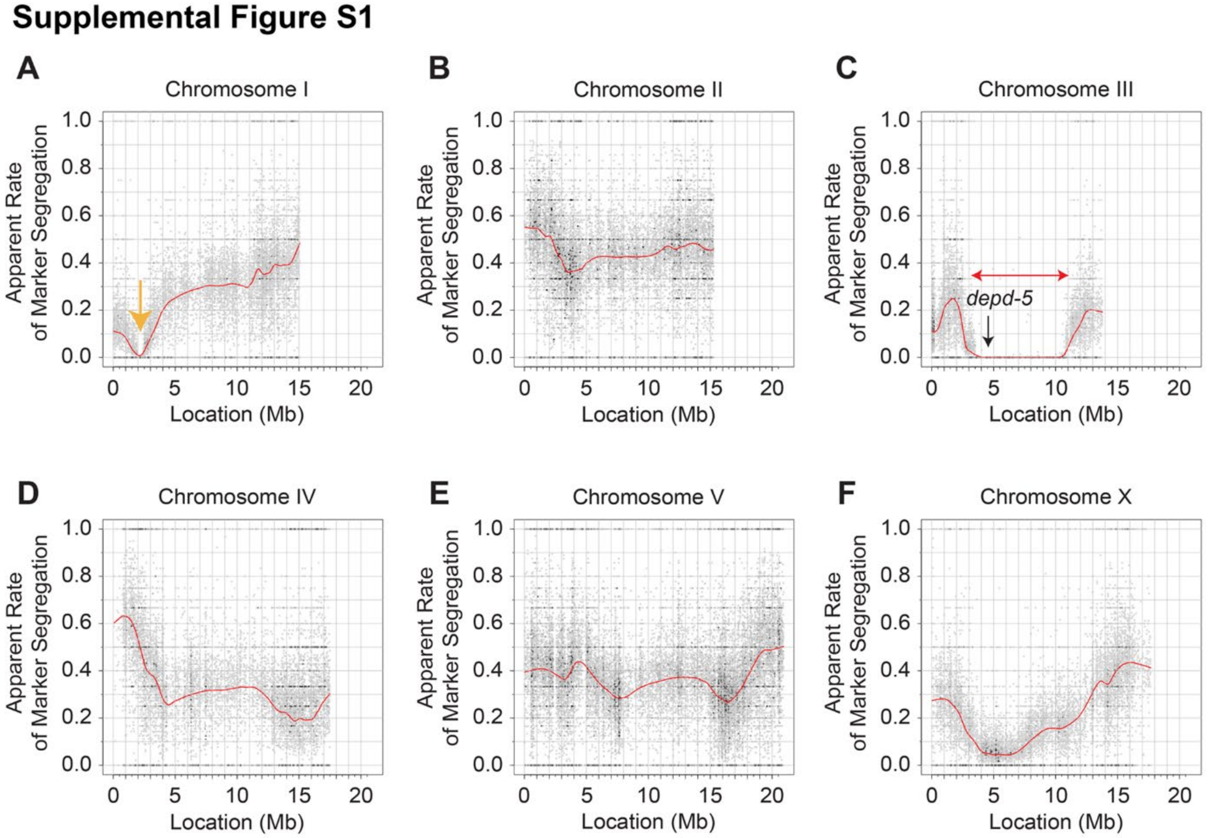

**Supplementary Figure S1. The *depd-5(am346)* mutation was positioned on Chromosome III by SNP mapping. (A-F)** To determine the position of the *depd-5(am346)* mutation in the *C. elegans* genome, we crossed *depd-5(am346); hlh-30(lf)* animals (Bristol N2 background) to the Hawaiian strain CB4856 with the *hlh-30(lf)* mutation introduced via CRISPR targeting and selected for animals homozygous for *depd-5(am346)*. This selection was performed several times, and then we used whole genome sequencing to detect SNPs from the CB4856 strain and newly arising EMS mutations. SNP-by-SNP allele frequency plots for Chromosomes I (A), II (B), III (C; also shown in Figure 1C), IV (D), V (E) and X (F) are shown. The horizontal axis shows the physical map of the chromosome (megabases), and the vertical axis shows the apparent rate of Hawaiian SNP marker segregation from 0 to 1.0, calculated as the number of CB4856 reads divided by the total reads at each SNP position. In panel C, the red horizontal arrow indicates the mapping region with a very low frequency of Hawaiian SNP markers and the black vertical arrow marks the approximate position of a candidate nonsense mutation in *depd-5*. In panel A, the yellow arrow depicts a region in chromosome I where a lack of Hawaiian SNP markers has been recurrently observed and is not specific to this experiment.

**Supplemental Figure S2**

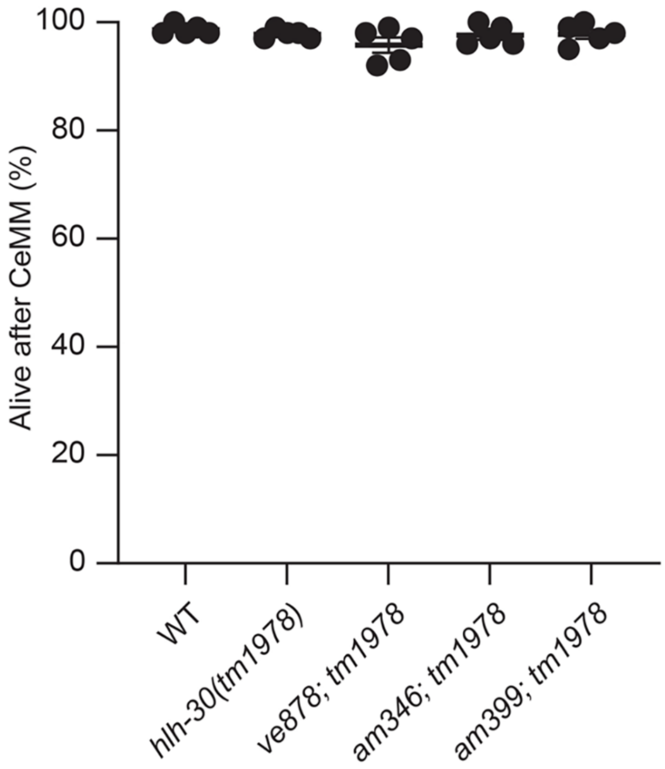

**Supplementary Figure S2. Glucose is not toxic to *depd-5(lf)* worms after CeMM exposure.** L1 larvae were starved for 33 hours, exposed to CeMM minimal with glucose and linoleic acid for 15 hours and scored for survival (following protocol shown in schematic in Figure 1B). The horizontal axis shows the genotypes as indicated and the vertical axis shows the percent of live animals. Bars and whiskers represent mean ± SEM (five independent experiments). No statistical significance was determined by ordinary one-way ANOVA.

#### Supplemental Figure S3

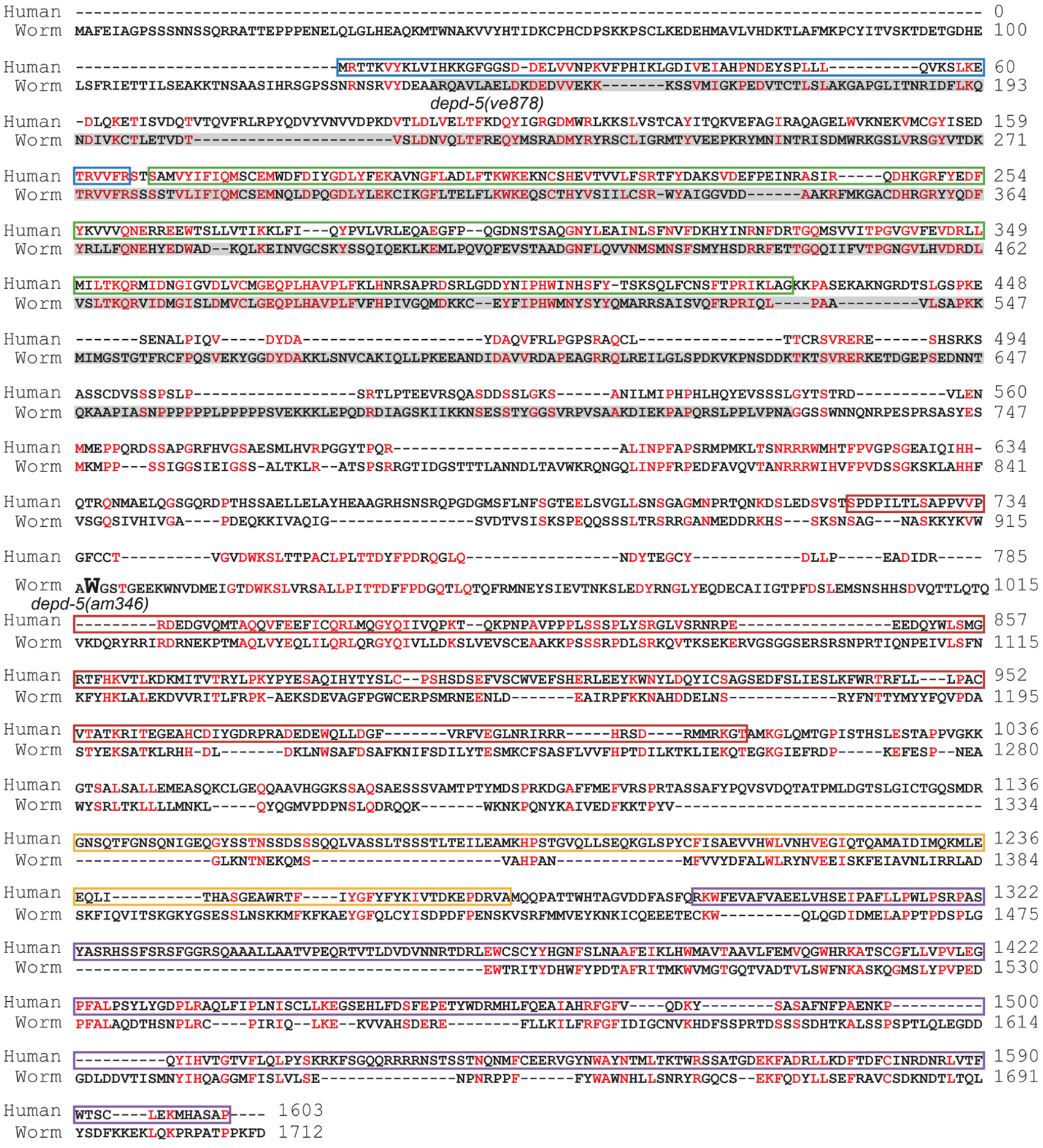

**Supplementary Figure S3. Amino acid alignment of human DEPDC5 and *C. elegans* DEPD-5.** Amino acid sequences of human DEPDC5 and *Caenorhabditis elegans* DEPD-5 were aligned using EMBL-EBI EMBOSS Needle to assess sequence conservation. Single-letter amino acid abbreviations are used: red amino acids are identical between human and worm. Numbers on the right indicate the last amino acid in each row. Conserved human DEPDC5 domains were defined according to Shen et al (see reference 14). The various domains are labeled as NTD (blue), SABA (green), SHEN (red), DEP (yellow), and CTD (purple). The *C. elegans depd-5(am346)* mutation causes a nonsense change, W917STOP, at the amino acid that is marked with a large font 'W'. The amino acids missing from the *depd-5(ve878)* deletion mutation strain are indicated by shading in grey.

### Supplemental Figure S4

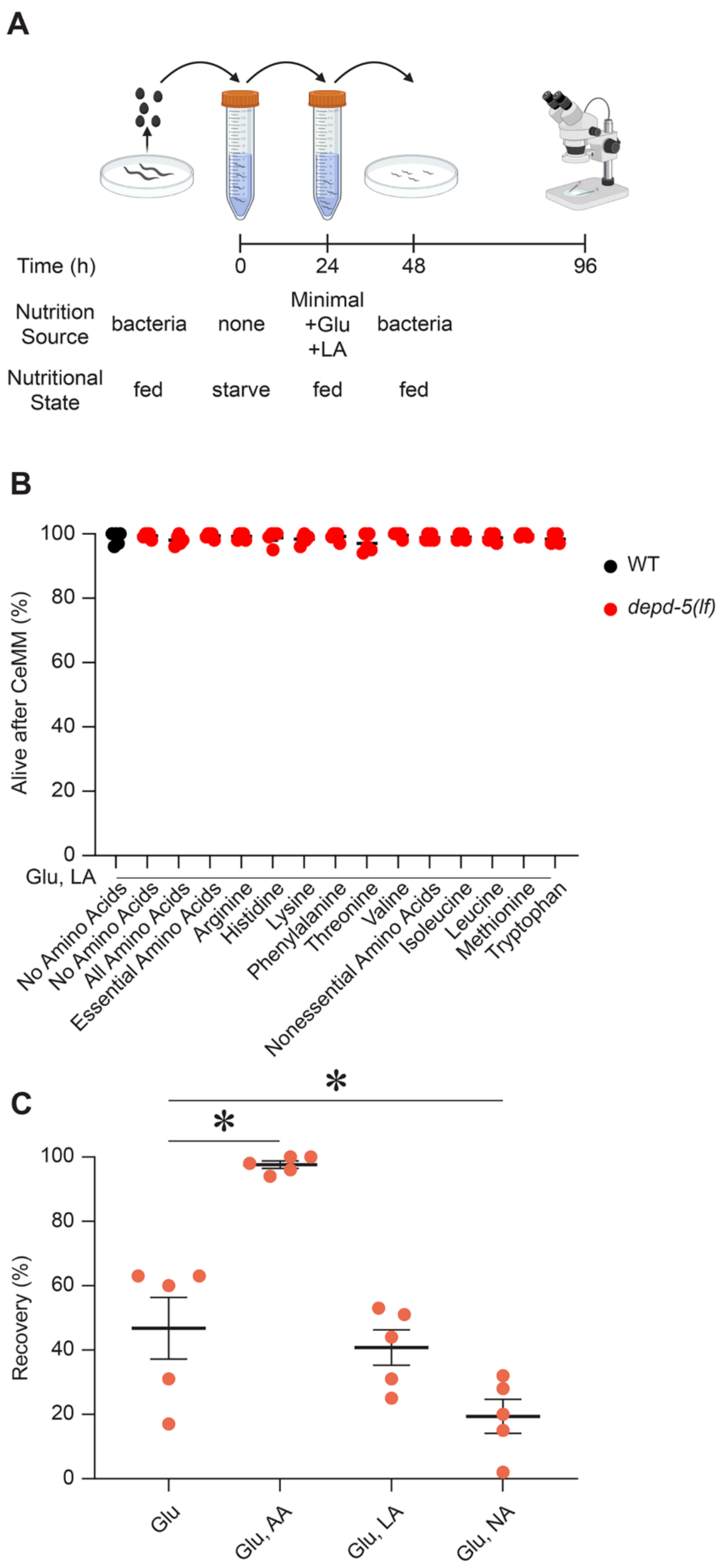

**Supplementary Figure S4. Amino acid supplementation does not affect survival during CeMM treatment following starvation. (A)** Experimental workflow for assessment of survival following starvation, refeeding with CeMM minimal and culture on *E. coli* overlaid on NGM agar plates. Animals were starved for 24 hours and then incubated for 24 hours in Minimal CeMM containing glucose and linoleic acid without or with amino acid or nucleic acid supplements (see B, C). **(B)** Wild type and *depd-5(lf)* L1 larvae were starved for 24 hours, exposed to CeMM minimal formulations supplemented with glucose, linoleic acid, and various amino acids as indicated for 24 hours, and scored immediately for survival (indicated as Alive after CeMM (%)). The horizontal axis shows the amino acids used and the vertical axis shows percent of animals alive after CeMM treatment of wild-type and *depd-5(am346)* animals. Bars and whiskers represent mean ± SEM (five independent experiments). Statistical significance among nutrient conditions was determined by ordinary one-way ANOVA. No significant differences were noted. **(C)** L1 larvae were starved for 24 hours, exposed to CeMM minimal formulations supplemented with glucose (Glu), glucose and amino acids (Glu, AA), glucose and linoleic acid (Glu, LA), or glucose and nucleic acids (Glu, NA) for 24 hours, cultured on *E. coli* OP50 layered agar dishes for 48 hours, and scored for survival (indicated as Recovery (%)). The horizontal axis shows the chemicals added to CeMM and the vertical axis shows recovery of *depd-5(am346)* animals. Bars and whiskers represent mean ± SEM (five independent experiments). Statistical significance was determined by ordinary one-way ANOVA followed by Dunnett's multiple-comparisons test comparing each treatment to the glucose-only group. $^*P < 0.05$.

### Supplementary Figure S5

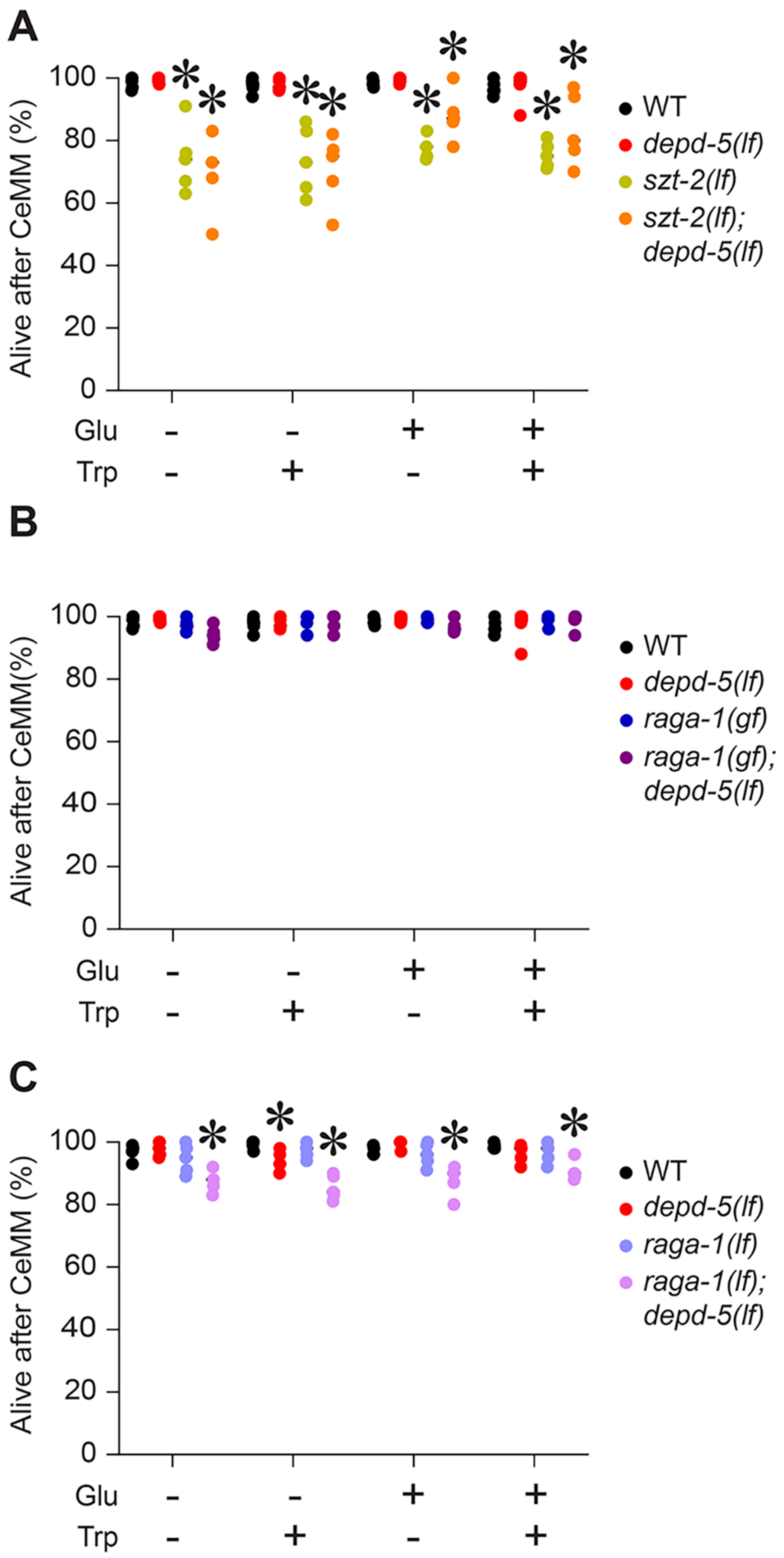

**Supplementary Figure S5. Survival of GATOR1 pathway mutants with exposure to nutrients following starvation. A-C)** L1 larvae of indicated genotypes were starved for 24 hours, exposed to CeMM minimal formulations supplemented with glucose (Glu), tryptophan (Trp), or both chemicals for 24 hours, and scored for survival. The horizontal axis shows the composition of CeMM and the vertical axis shows percent of animals that are alive following CeMM treatment. Bars and whiskers represent mean ± SEM (five independent experiments). Percent recovery data were transformed using an arcsine square-root transformation before statistical analysis. Transformed values were analyzed by ordinary two-way ANOVA followed by Tukey's multiple-comparisons test. *$P < 0.05$ vs. WT control within the respective nutrient group.

#### Supplementary Figure S6

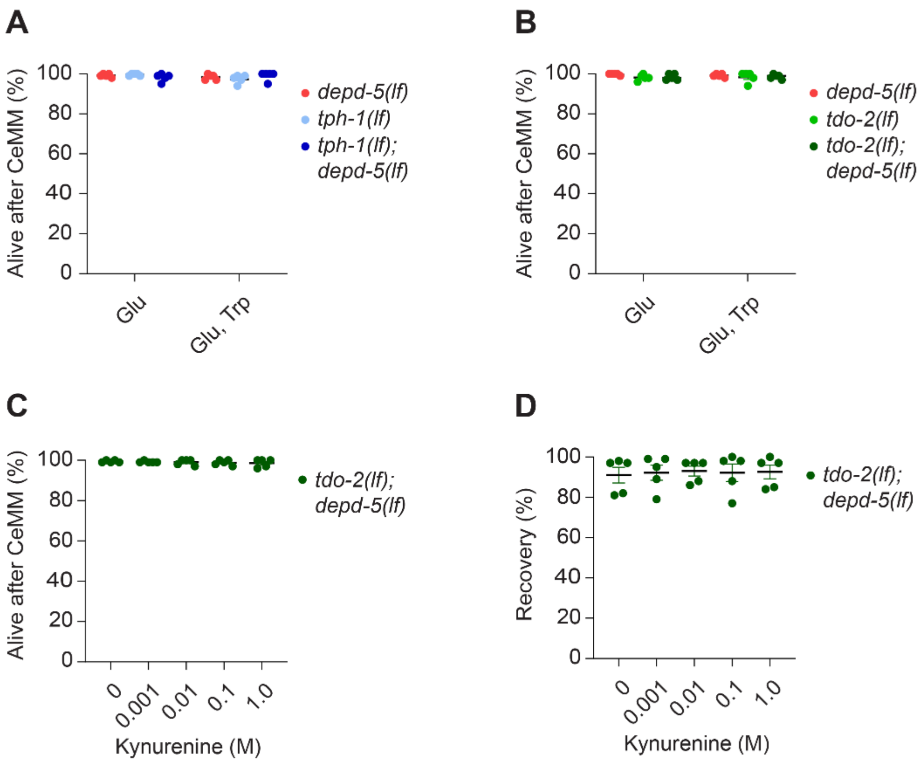

**Supplementary Figure S6. Survival of tryptophan catabolism mutants with exposure to nutrients following starvation. (A, B)** L1 larvae of worm strains with the indicated genotypes were starved for 24 hours, exposed to CeMM minimal formulations supplemented with glucose (Glu) or glucose and tryptophan (Glu, Trp) for 24 hours, and immediately scored for survival (assessed as Alive after CeMM (%)). The horizontal axis shows the composition of CeMM minimal and the vertical axis shows percent of animals that are alive after the CeMM treatment. Bars and whiskers represent mean ± SEM (five independent experiments). Percentage survival data were transformed using an arcsine square-root transformation before statistical analysis. Transformed values were analyzed by ordinary two-way ANOVA followed by Šídák's multiple-comparisons test comparing genotypes within each nutritional condition. No statistically significant differences were noted. **(C, D)** L1 larvae of *tdo-2(lf); depd-5(lf)* double mutant animals were starved for 24 hours and exposed to CeMM Minimal formulations supplemented with increasing concentrations of kynurenine (0 - 1.0 M) for 24 hours. (C) Animals were scored for survival immediately following CeMM exposure (assessed as Alive after CeMM (%)). (D) Animals surviving CeMM exposure were scored for recovery after 48 hours on OP50-seeded NGM dishes (assessed as Recovery (%)). The horizontal axis shows kynurenine concentration and the vertical axis shows percent survival after CeMM (C) or percent recovery (D). Symbols represent individual experiments and the horizontal bars represent mean ± SEM (five independent experiments). Data were analyzed by ordinary one-way ANOVA followed by Dunnett's multiple comparisons test comparing each kynurenine concentration with the untreated control. No significant differences were detected in survival or recovery.

### Supplementary Figure S7

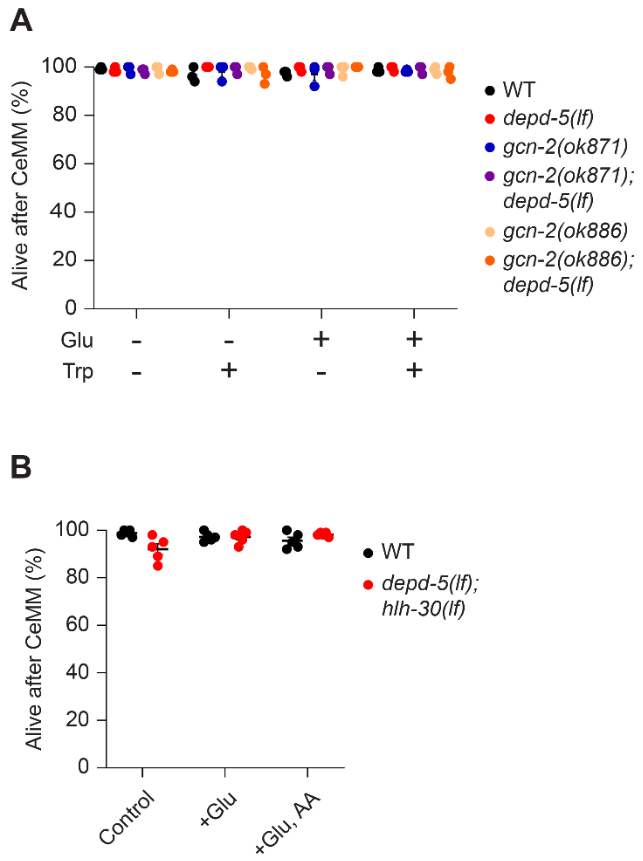

**Supplementary Figure S7. Survival followed starvation and exposure to CeMM formulations in nutrient-signaling mutants. (A)** L1 larvae of worm strains with the indicated genotypes with *gcn-2(lf)* with and without *depd-5(lf)* and wild type controls were starved for 24 hours, exposed to CeMM minimal formulations supplemented with glucose (Glu) or glucose and tryptophan (Glu, Trp) for 24 hours, and immediately scored for survival (assessed as Alive after CeMM (%)). The horizontal axis shows the composition of CeMM minimal and the vertical axis shows percent of animals that are alive after the CeMM treatment. Bars and whiskers represent mean ± SEM (three independent experiments). Percentage survival data were transformed using an arcsine square-root transformation before statistical analysis. Transformed values were analyzed by ordinary two-way ANOVA followed by Tukey's multiple-comparisons test comparing genotypes within each nutritional condition. **(B)** L1 larvae of worm strains with the indicated genotypes with *depd-5(lf)*;*hlh-30(lf)* double mutants and wild type controls were starved for 24 hours, exposed to CeMM minimal formulations supplemented with glucose (Glu) or glucose and tryptophan (Glu, Trp) for 24 hours, and immediately scored for survival (assessed as Alive after CeMM (%)). The horizontal axis shows the composition of CeMM minimal and the vertical axis shows percent of animals that are alive after the CeMM treatment. Symbols represent individual experiments and the horizontal bars represent mean ± SEM (five independent experiments). Percentage survival data were transformed using an arcsine square-root transformation before statistical analysis. Transformed values were analyzed by ordinary two-way ANOVA followed by Tukey's multiple-comparisons test comparing genotypes within each nutritional condition. Only comparisons with wild type within each nutritional condition are indicated. Data are from five independent experiments. $*P < 0.05$.

### Supplementary Figure S8

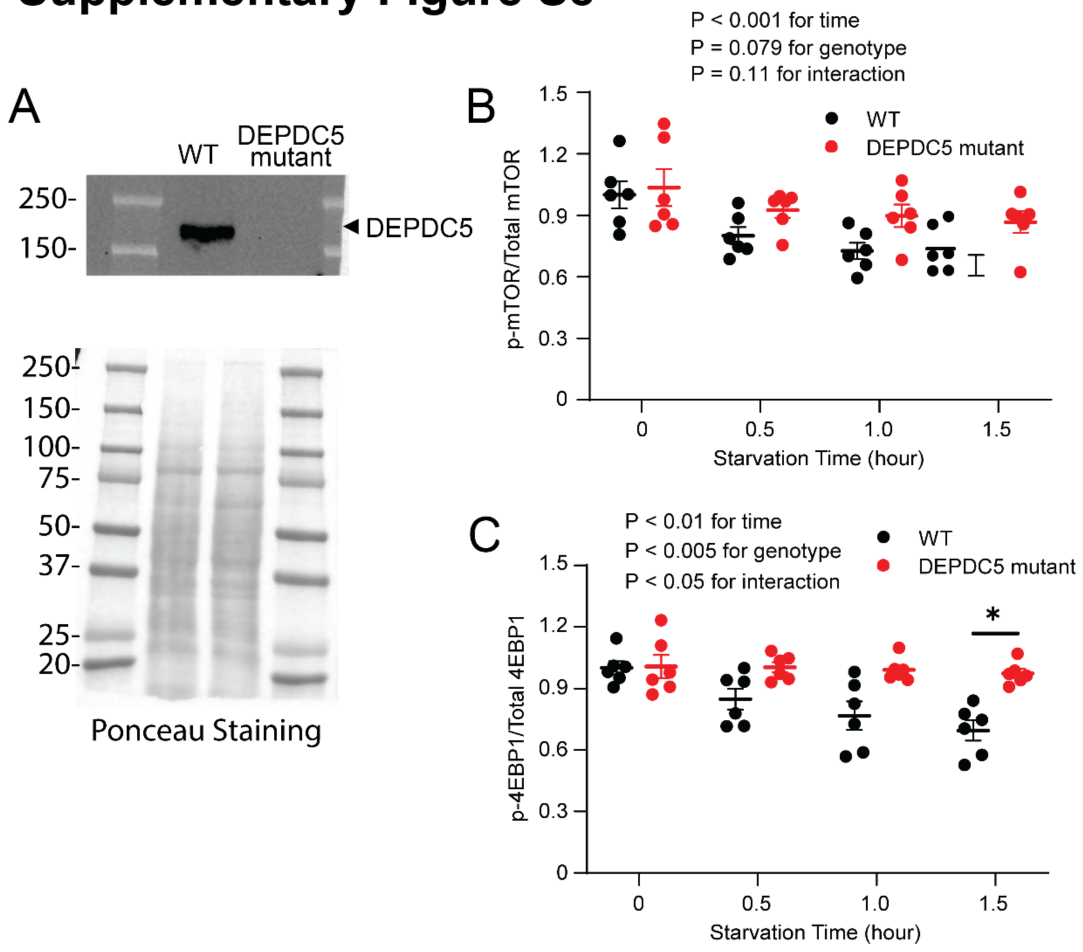

**Supplementary Figure S8. TORC1 signaling during starvation in DEPDC5 mutant HeLa Cells. (A)** Immunoblot analysis of DEPDC5 in wild-type and DEPDC5 mutant HeLa cells. Ponceau S staining for total proteins is shown as loading control. **(B-C)** Quantification of (B) phosphorylated mTOR relative to total mTOR and (C) phosphorylated 4EBP1 relative to total 4EBP1 in wild type and DEPDC5-KO HeLa cells following the indicated durations of starvation. Data are expressed as fold over WT at t=0 (six independent experiments). Statistical significance was determined by ordinary two-way ANOVA followed by Tukey's multiple-comparisons test. **P*<0.05.
